# Mapping genetic effects on cellular phenotypes with “cell villages”

**DOI:** 10.1101/2020.06.29.174383

**Authors:** Jana M. Mitchell, James Nemesh, Sulagna Ghosh, Robert E. Handsaker, Curtis J. Mello, Daniel Meyer, Kavya Raghunathan, Heather de Rivera, Matt Tegtmeyer, Derek Hawes, Anna Neumann, Ralda Nehme, Kevin Eggan, Steven A. McCarroll

## Abstract

Tens of thousands of genetic variants shape human phenotypes, mostly by unknown cellular mechanisms. Here we describe Census-seq, a way to measure cellular phenotypes in cells from many people simultaneously. Analogous to pooled CRISPR screens but for natural variation, Census-seq associates cellular phenotypes to donors’ genotypes by quantifying the presence of each donor’s DNA in cell “villages” before and after sorting or selection for cellular traits of interest. Census-seq enables population-scale cell-biological phenotyping with low cost and high internal control. We demonstrate Census-seq through investigation of genetic effects on the SMN protein whose deficiency underlies spinal muscular atrophy (SMA). Census-seq quantified and mapped effects of many common alleles on SMN protein levels and response to SMN-targeted therapeutics, including a common, cryptic non-responder allele. We provide tools enabling population-scale cell experiments and explain how Census-seq can be used to map genetic effects on diverse cell phenotypes.

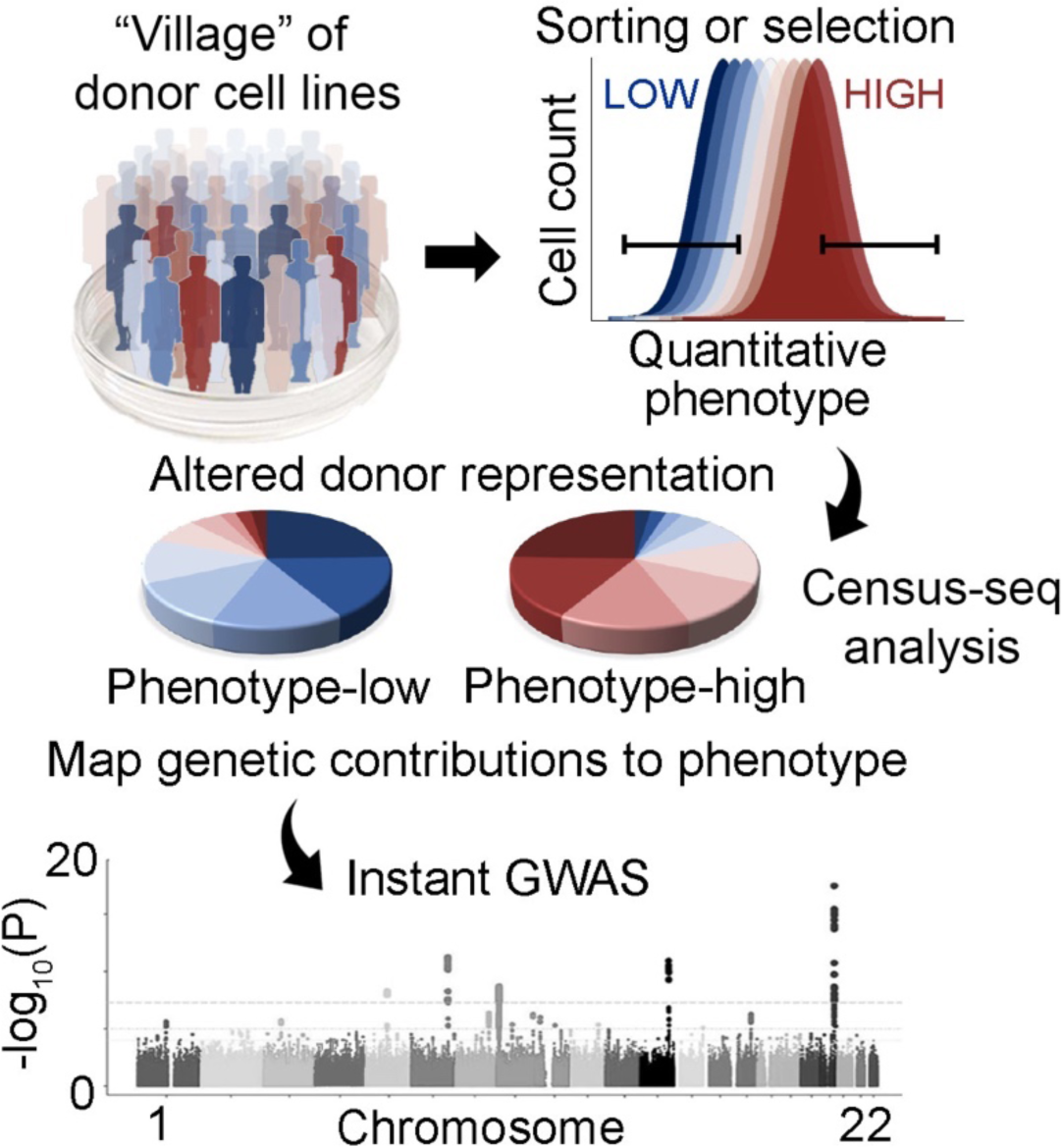

**Highlights:**

- Census-seq reveals how inherited genetic variation affects cell phenotypes
- Genetic analysis of cellular traits in cell villages of >100 donors
- Characterizing human alleles that shape SMN protein expression and drug responses
- Development of protocols and software to enable cellular population genetics

## Introduction

Human populations harbor vast numbers of common and rare alleles; such alleles affect the protein-coding sequence or regulation of almost all human genes. Human genetic studies have associated tens of thousands of alleles to risk of illnesses and other quantitative traits. A core goal of human genetics is to help identify cellular processes that underlie disease. And yet we know little today about how human alleles affect cells and their biology. We understand even less about how combinations of alleles – whether from one or many genes – converge upon cell-biological processes that might mediate normal variation and vulnerabilities.

Pioneering studies have shown that lymphoblastoid cell lines or pluripotent stem cells (PSCs) from human donors can be used to identify how common DNA variation shapes certain cellular phenotypes, especially RNA expression (Cheung et al., 2005, Morley et al., 2004, Stranger et al., 2007a, Stranger et al., 2007b, Kilpinen et al., 2017, Lo Sardo et al., 2017, McFarland et al., 2019, Pickrell et al., 2010). However, efforts to ascertain effects of human genetic variation on cellular phenotypes encounter two challenges. The first challenge involves reaching the necessary scale by culturing and assaying the large number of cell lines necessary to associate phenotype with genotype. Challenges of scale have largely limited genome-wide genetic studies to a few labs or consortia and a few phenotypes. The second challenge involves control and rigor: how to accurately measure and compare phenotypes across many cell lines cultured separately. Without such control, it is often feared that biology can learn only from “alleles of large effect” – alleles that cause dramatic phenotypes in deterministic ways and thereby overwhelm noise in experimental measurement.

Here we describe an experimental system and computational methods (“Census-seq”) that we developed to perform population-scale cellular experiments that enable insights from genetic influences of all kinds and frequencies upon diverse cellular phenotypes. Our approach involves what we call “village-in-a-dish” experiments, in which cells from all donors are mixed together, then fed, passaged, and stimulated in a shared environment, and finally scored for phenotypes all together. Census-seq analysis relates cells’ phenotypes to the individual cell donors by analyzing the donors’ DNA contributions to cell mixtures: after sorting or selecting the cell village for the phenotype of interest, we sequence the genomic DNA in the resulting, derived villages; computational analysis reveals the proportion of cells from each donor before and after sorting or selection. This approach allows many different kinds of cellular phenotypes to be analyzed for association to donor genotypes or other donor characteristics.

Census-seq addresses many challenges that have limited cellular genetic studies. Our first goal has been to make it facile and inexpensive to do population-scale phenotype readouts and genetic analysis with cells. A second goal has been to measure phenotypes across scores of cell lines in a rigorous, well-controlled way. A third goal has been to facilitate genetic analysis of phenotypes beyond RNA expression: much information flows across cellular networks through proteins, which may provide a natural integration point for effects from many genes and alleles. Cell villages dramatically reduced the cost and complexity of population-scale experiments while reducing measurement noise and biological variance.

We sought here to establish such systems, understand their practical execution, apply them to a model phenotype, and enable other scientists to adopt similar approaches. We chose as a model phenotype the expression of the SMN protein, for which deficiency underlies Spinal Muscular Atrophy (SMA), a common congenital disorder engaged by emerging therapeutics (Chen, 2020, Lefebvre et al., 1995).

## Results

### Census-seq: determining the donor composition of a cellular “village”

Census-seq is analogous to pooled CRISPR screening, with a key difference: Census-seq interrogates natural genetic variation rather than synthesized libraries of guide RNAs. In pooled CRISPR screens, cells are administered a library of gene-perturbing guide RNAs (each tagged with a DNA barcode) and then sorted or selected for a phenotype of interest; the relative frequencies of barcodes are compared between the initial population of cells and the population created by selection or sorting, and the effects of each guide on the phenotype are inferred from the change in that guide’s representation (Adelmann et al., 2019, Canver et al., 2015, Gasperini et al., 2019, Hsu et al., 2018, Shalem et al., 2014, Wang et al., 2014). For Census-seq, we begin by constructing villages of cell lines derived from many (most frequently 10-100) donors; the cells in the village may be cultured or stimulated together over some period of time. Then, as in pooled CRISPR screening, we fractionate the cell village by sorting (or performing selections) to obtain cells with a phenotype of interest. By sequencing genomic DNA derived from the cell village, and applying a computational approach we describe below, we then determine the relative contribution of each donor’s genomic DNA to the cell mixture – and therefore the fraction of all cells that come from each donor – comparing the initial population to the population that results from fractionation, or comparing fractionated populations to one another.

Census-seq uses natural genetic variation (rather than synthetic barcodes (Yu et al., 2016)) to measure each individual’s contribution to cell mixtures. The use of inherited variation as a natural barcode makes it possible to use human cells without the further perturbations (e.g. viral transfection and cloning) that alter cells’ biology or contribute to the acquisition of mutations. To infer each donor’s cellular contribution to a mixture, we isolate genomic DNA from the village, then perform low-coverage whole genome sequencing on that genomic DNA (generally about 1X average genomic coverage, at a cost of about $100 per cell village). We routinely analyze as few as 1×10^5^ cells of a village, enabling many experiments to be performed in relatively small reaction chambers such as those on a 12-well plate.

We developed a computational approach to estimate each donor’s contribution to this DNA mixture (**Figure 1A**). Making this estimate requires *a priori* genetic information on the individual donors, which can come from SNP arrays, exome sequencing, or whole genome sequencing (WGS) (**Figure S1E**). Sequencing the village’s genomic DNA generates millions of sequence reads; these reads sample the donors’ genomes in proportions that reflect the representation of each donor’s cells in the village. A large minority of the sequence reads are “allelically informative” in the sense that they contain a genomic site that we know to vary among the potential donors; for example, in an analysis of a village of 40 donors whose genetic variation has been ascertained by WGS, about 42% of all 151-bp reads contain a genomic site for which the donors have varying genotypes. The allele present on any such read offers partial information about the composition of the mixture, as only a subset of the potential donors’ genomes can be sources of that sequence read.

**Figure 1.**
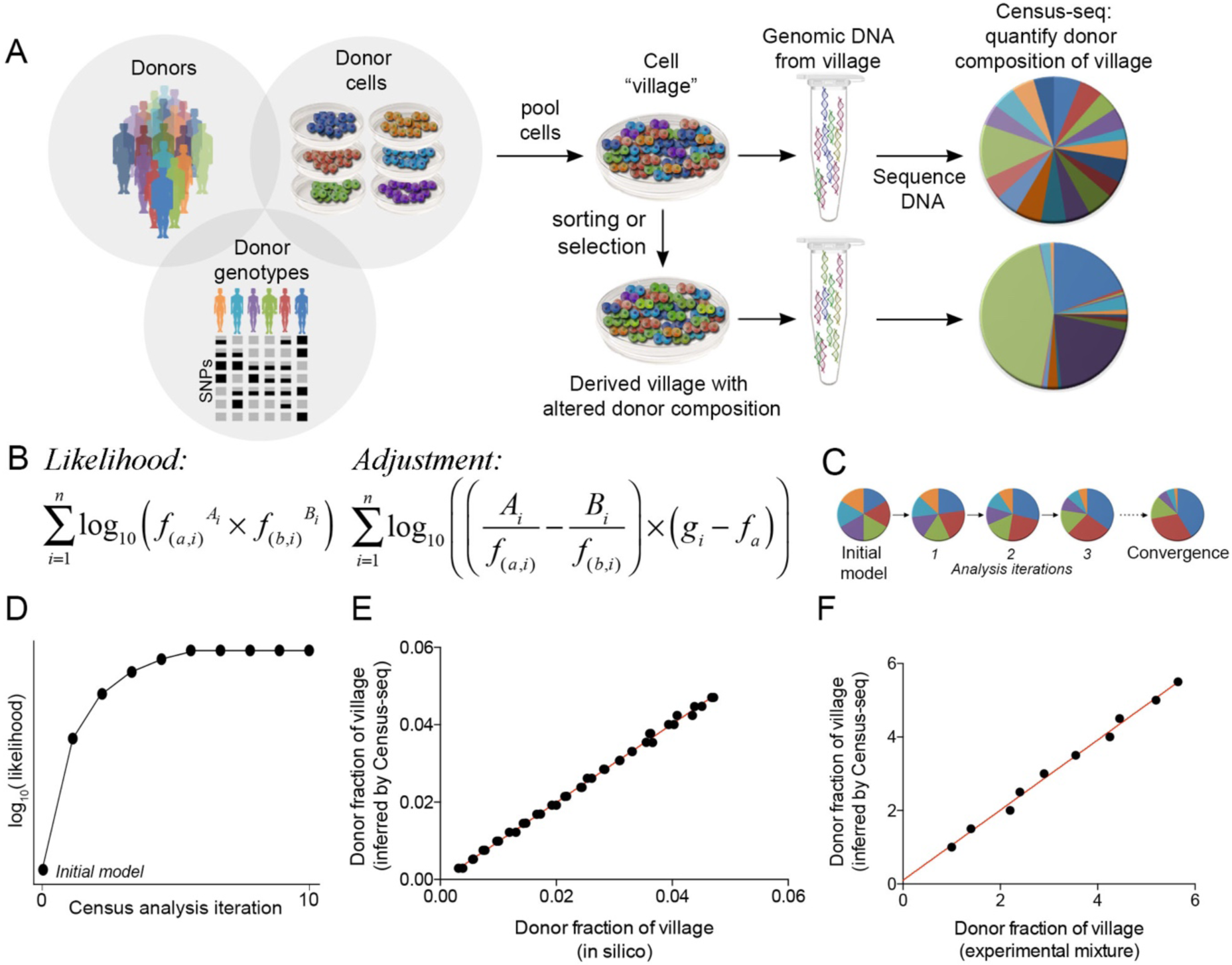
Village-in-a-dish experimental systems. A. Cells from many donors are cultured together as a “village” to enable scalability and minimize technical sources of variation. Sorting or selection enrich for cells with phenotypes of interest, creating a derived village. Genomic DNA is extracted from each village and sequenced. Census-seq analysis measures each donor’s contribution to the village’s genomic DNA, and thus (indirectly) the relative number of cells from each donor in each village. B. Census-seq analysis is based on an expectation-maximization (EM) algorithm. The algorithm seeks the set of donor-mixing coefficients (summing to 100%) that maximize the likelihood of the observed sequence data. For any set of coefficients, this likelihood is measured by multiplying the modeled allele frequencies (in the village mixture) of all of the alleles observed on all allelically informative sequence reads. The donor-mixing coefficients are then adjusted in the direction that most strongly increases the likelihood of the observed data; the adjustments are derived by taking the derivative of the likelihood with respect to each of the donor-specific mixing coefficients (Methods) (*f*_*a*_ = the frequency of the reference allele, *A* = the counts of the reference allele in the sequence data, *f*_*b*_ = the frequency of the alternate allele, *B* = the counts of the alternate allele in the sequence data, *g*_*i*_ = the genotype of the donor at the site, formatted as 1 for the reference genotype, 0.5 for the heterozygous genotype, and 0 for the alternate allele) C. Iterated rounds of adjustment optimize the estimates of donor mixing coefficients to fit the sequencing data, converging asymptotically to a final estimate. D. Convergence is typically reached after just a few iterations of the EM algorithm. E. Simulated “village DNA” data sets were made by mixing whole genome sequence data from 40 unrelated donors in such a way that each donor contributed a different proportion of the overall data. Census-seq was used to estimate the quantitative contribution of each donor to this data mixture. The plot compares the known, *in silico* mixing coefficients to the estimates from Census-seq. F. Genomic DNA from ten donors was mixed in such a way that each donor contributed a different proportion of the total DNA. The DNA mixture was sequenced and analyzed by Census-seq. The plot compares the aliquoted donor-contribution proportions to the Census-seq estimates from sequencing the DNA mixture.

In the mathematical analysis underlying Census-seq, we find the set of mixing coefficients (one coefficient for each donor, summing to 1.0) that make the observed sequence data – the millions of allelically informative reads, considered together – maximally likely to have been generated by random sampling from the donors’ genomes (**Figure 1B**). The mixing coefficients are inferred via an expectation-maximization algorithm which works in the following way. At every variable site in the human genome, any hypothetical mixture of the donors involves an implicit “village allele frequency” for every allele in the DNA mixture (**Figure 1B**). (A default initial condition for analysis can be that each donor has contributed equal numbers of cells; in this case the village allele frequency for each allele is simply that allele’s frequency among the donors, without weighting.) We measure the likelihood of the village’s sequence data by multiplying the village allele frequencies of all of the alleles from allelically informative sequence reads, making small adjustments to account for the possibility of sequencing error. To refine the mixing coefficients, we calculate the partial derivative of the likelihood of the data with respect to each donor’s mixing coefficient; this calculation yields a set of adjustment factors by which we increase or decrease each individual donor’s mixing coefficient to improve the data likelihood (**Figure 1B, C, D, Methods**). This “gradient ascent” process is repeated; as the mixing coefficients are adjusted, the likelihood of the observed sequence data increases toward an asymptote (**Figure 1D**). The computational analysis converges quickly – typically requiring just 10-30 iterations – to a set of mixing coefficients under which the observed sequence data are as likely as possible.

To evaluate whether the donor composition of villages inferred by Census-seq corresponded to their known, actual composition, we performed many control analyses and experiments, including (i) analyzing *in silico* simulations in which we mixed DNA sequencing data from many individuals in known proportions; (ii) mixing genomic DNA from different individuals in known concentrations; and (iii) mixing cells from individuals in known proportions. In each case, the Census-seq estimates of donor representation in the mixture corresponded closely to the ratios in which sequence data, DNA or cells from different donors had been mixed (**Figure 1E, F, S1**). Overall, we found that a donor’s DNA representation in a village could be measured accurately down to a limit at which a donor contributes about 0.2% of the cells in a mixture (**Figure S1F**), a limit that is related to the sequencing error rate and thus is not addressed by simply sequencing Census-seq libraries more deeply. This limit places an upper bound (of a few hundred) on the number of unique donors that can be accurately quantified in one village. This bound is still far above the scale of experiments that can be accomplished comfortably in traditional formats; still-larger experiments are possible by meta-analyzing many villages with overlapping membership for calibration.

We found that donor composition of 40-donor villages could be inferred accurately with modest amounts of sequencing that corresponded to less than 1X coverage of a single human genome (**Figure S1E)**. The required depth of sequencing depended on the complexity of the village (number of donors) and the amount of available genome information on these donors (**Figure S1E**) (**Methods**), with deeper *a priori* genetic characterization (e.g. WGS) causing more sites to be allelically informative and thus allowing lighter sequencing of the village’s DNA at the time of phenotype analysis. For example, if a donor has contributed 2.0% of the DNA in a mixture, sequencing the village genomic DNA to a sequencing depth of about 1X (16 million 150-bp reads) yields estimates of 2.0 +/- 0.1%. We routinely analyze 16 villages in each run of a desktop sequencer (Illumina NextSeq) (**Methods**) at a cost of about $100 per village.

These results encouraged us to use this approach to analyze a great many cell villages of experimental composition and to study the population dynamics of villages of PSCs as they grew in culture together. While PSCs present important opportunities in their capacity to differentiate *in vitro* into a great many cell types, PSCs present particular challenges for cell villages, as they are proliferative and acquire mutations in culture. The results were sobering.

### Population dynamics of cellular villages

To study the population dynamics of PSC villages, we established 29 villages, each consisting of ∼10-100 donors, and measured their population dynamics across 3-13 passages; in total, we measured 3,705 Census-seq growth phenotypes in cell lines from 247 unique donors at multiple timepoints (**Figure 2A-E, Table S1, S2**). For initial experiments we utilized cell lines drawn from a large collection of human embryonic stem cells we had assembled and recently genetically characterized by whole genome sequencing (Merkle et al., in revision, Merkle et al., 2017). In most experiments, we found that one or a few cell lines progressively took over the village (**Figure 2C, D**). Analyses of the whole genome sequences of these cell lines indicated that a majority of the hyperproliferative cell lines had acquired growth-promoting mutations (**Figure 2C**), some of which – such as mutations in the *TP53* gene and the gene encoding the p53 inhibitor MDM4 – recurred in multiple cell lines (Loh et al., 2018, Merkle et al., 2017). The ease and low cost of Census-seq analyses made it straightforward to detect cell lines with these growth promoting mutations (**Figure 2C, D**). Though these findings demonstrated the utility of Census-seq for identifying stem cell lines with growth-promoting mutations (that would ideally be excluded from translational efforts), they also indicated that variation in growth rates among stem cell lines must be managed to maintain large villages.

**Figure 2.**
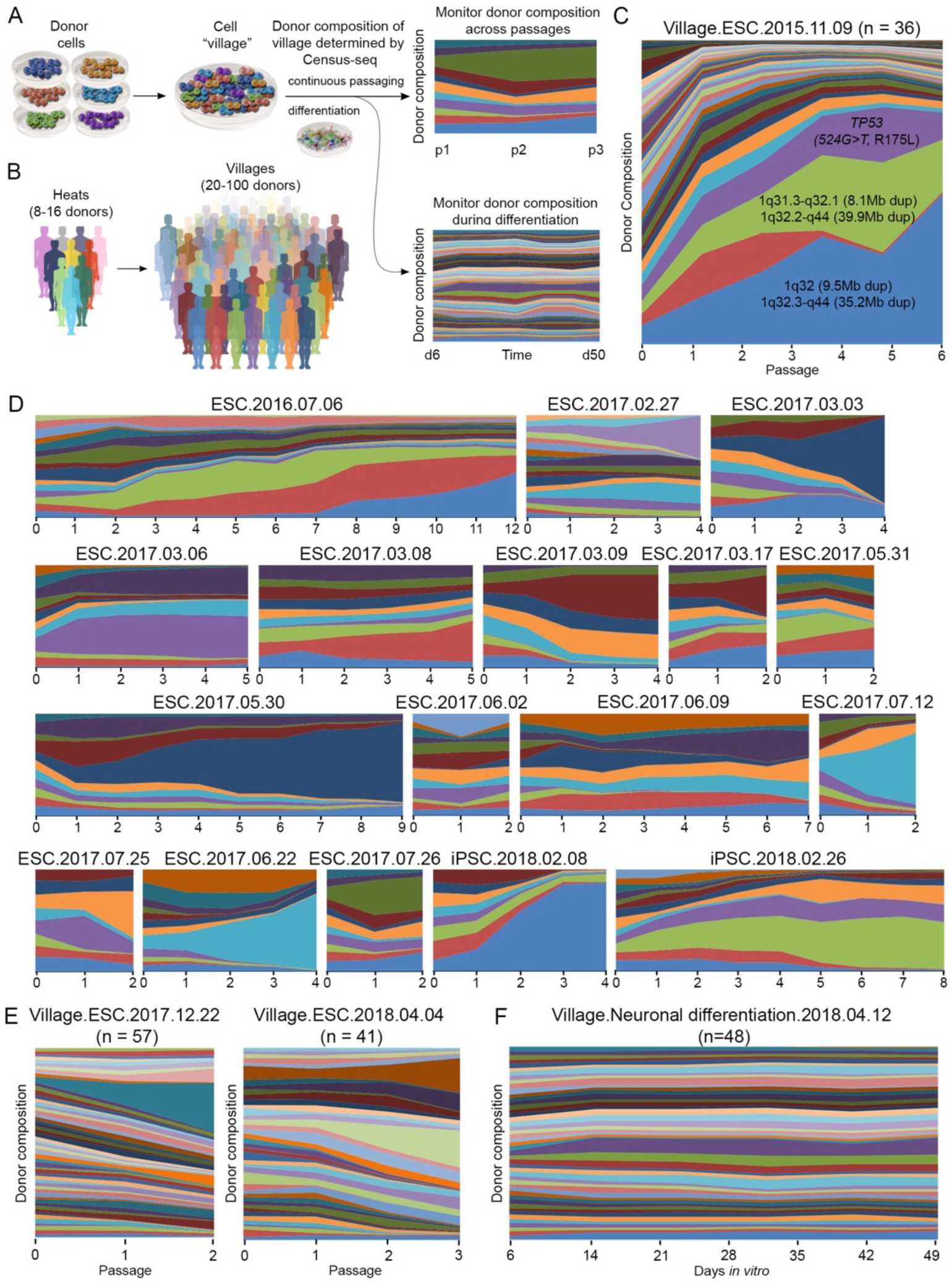
Donor population dynamics in villages of pluripotent stem cells. A. Villages were created from PSCs and cultured or differentiated *in vitro*. Census-seq was used to monitor the donor composition of these cultures at each passage. B. Before creating large villages, smaller “heats” can be used to identify cell lines that are hyper-proliferative. C. Several hyper-proliferative PSC lines had acquired growth-promoting mutations in culture. D. Hyper-proliferative lines can quickly distort the donor composition of villages, rendering them unsuitable for many kinds of experiments. E. A variety of steps, including excluding lines with recognizably culpable acquired mutations and other empirically hyper-proliferative lines, allow villages to be maintained across several passages with relatively stable donor composition. F. Villages of PSCs can be differentiated into neurons and cultured for weeks as the neurons mature *in vitro*, with stable donor composition of the village.

We therefore used Census-seq to explore approaches that would allow us to better scale the size of villages. Overall, we found that constructing larger villages of 40-100 cell lines was more robust if we first established and then monitored the population dynamics of small “heats” of 8-12 PSC lines (**Figure 2D**). Census-seq analysis of these heats made it possible to identify cell lines that were empirically hyper-proliferative, regardless of whether they contained obviously culpable acquired mutations, before then choosing which cell lines to include in larger villages. For instance, in 7 early heats in which we evaluated 67 cell lines, we identified 13 cell lines (19%) that exhibited rapid expansion relative to the other lines (**Table S3**). Eliminating these rapidly growing outliers greatly increased the compatibility of cell lines within a village, allowing the maintenance of donor balance over longer co-cultures that were amenable to numerous forms of experimental manipulation and cellular phenotyping (**Figure 2E**).

We next assessed the dynamics of villages during directed stem cell differentiation, which makes it possible to investigate genetic influences on phenotypes found only in specific cell types. As an initial test, we employed a rapid, reproducible neuronal differentiation strategy to generate homogenous pools of post-mitotic neurons by inducing the expression of Neurogenin 2 (*Ngn2*) combined with small-molecule patterning of cells towards a cortical excitatory identity (Nehme et al., 2018, Zhang et al., 2013). This approach generated neuronal villages that matured over many weeks while maintaining a stable mixture of the donors (**Figure 2F**), useful for perturbation and phenotypic profiling in downstream assays.

### A computational toolkit for village experiments: “Roll Call” and “CSI”

Simulations showed that the precision of Census-seq analyses would be undermined if a cell village was contaminated with cells from a donor who was not expected to be present in a village. To address this, we developed computational approaches to quickly authenticate a village (or other DNA mixture) and identify and diagnose the presence of cells with unexpected genomes (**Figure 3, Methods**). These methods have the added benefit of being generally useful for validating the provenance of large numbers of cell lines (Nelson-Rees et al., 1981).

**Figure 3.**
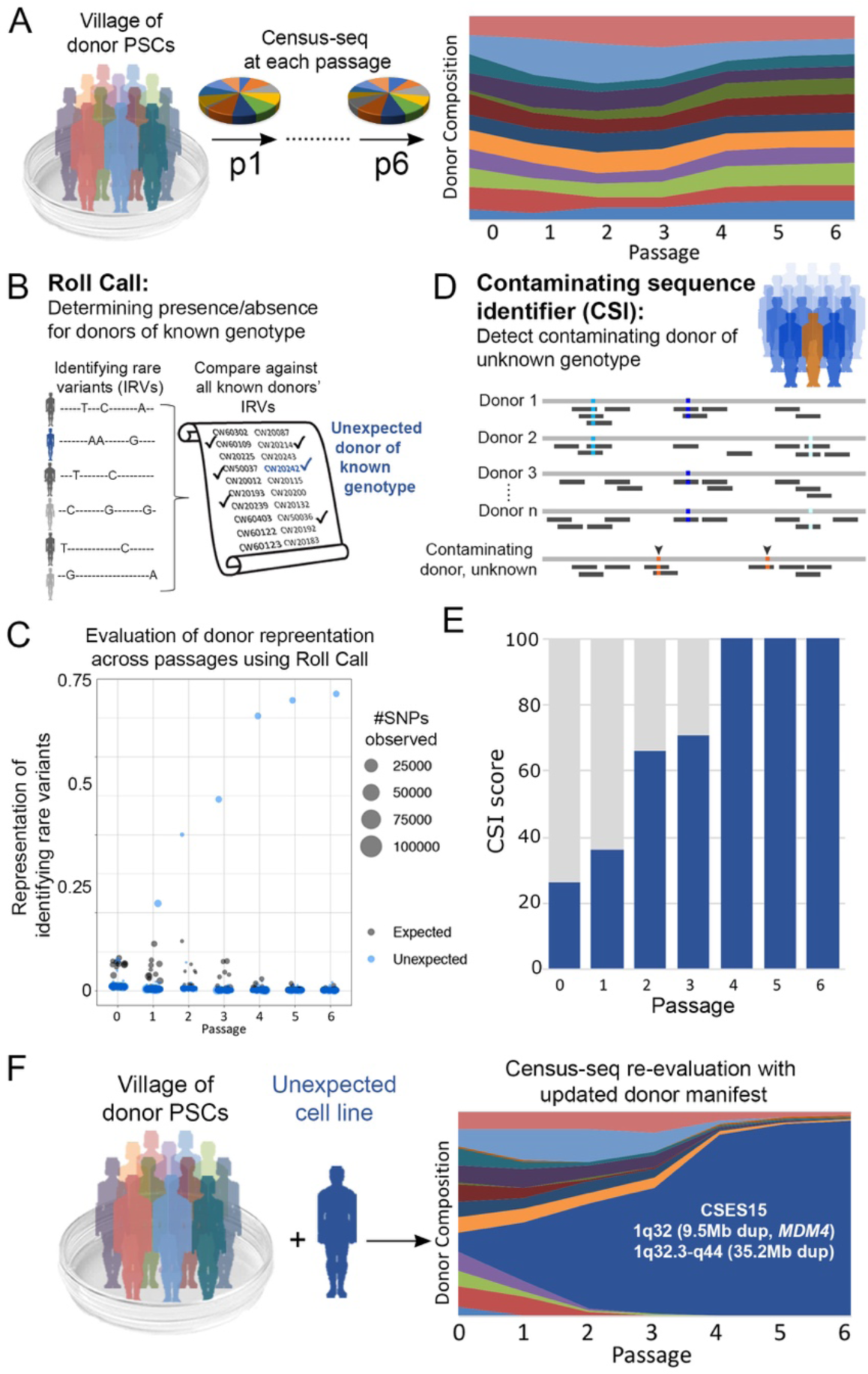
Using Roll Call and CSI to inspect donor composition of villages. A. In a typical analysis, Census-seq is used to determine donor composition of a village at each passage. The appearance of a strikingly balanced village after 6 weeks in culture prompted the generation of additional analysis tools to further scrutinize the donor composition of this and other cell villages. B. The Roll Call algorithm is used to confirm the presence or absence of individual donors (with known, *a priori* genetic variation information). Roll Call utilizes Identifying Rare Variants (IRVs) – rare variants that distinguish individual candidate donors from all other candidates. C. Roll Call identified the intrusion of a familiar (but unexpected) cell line into a village, indicating that this line (light blue) dominated donor representation within a few passages. D. The Contaminating Sequence Identifier (CSI) algorithm detects the presence of DNA from unexpected donors of unknown genotype (orange). E. In an *in silico* analysis blinded to any genetic information about the contaminating cell line, CSI predicted that an unfamiliar donor was present and had increased representation with each passage. F. Re-evaluating the donor composition of this village using Census-seq with an updated roster of donor cell lines confirmed the presence on a dominant intruding cell line. This line was determined to have acquired multiple growth-promoting mutations, including within *MDM4*, which encodes a regulator of the P53 tumor suppressor.

Contamination of a village could in principle arise from a donor of known genotype, or a donor of unknown genotype; we developed two computational approaches to address these two cases. To illustrate these approaches, we have drawn on the example of a village whose 12 donors initially appeared to have maintained a remarkably stable balance through six passages (**Figure 3A**).

To detect the presence of donors with known genome sequences, we developed “Roll Call”, which utilizes variants that distinguish individual donors from all other donors for whom cells might be present in a given lab or project (**Figure 3B, S2A-D**); we call such variants Identifying Rare Variants (IRVs). The presence of a sufficient number of sequence reads with an individual donor’s IRVs confirms the presence of that donor’s cells in a village. In the experiment in question, Roll Call identified that the village DNA contained a great many IRVs from an unexpected cell line (CSES15) – a cell line whose genome we had previously analyzed, but was not meant to be included in the village – and suggested that this cell line had become more abundant in the village with each passage (**Figure 3C**). Identifying the contaminating donor made it possible to correct the Census-seq analysis to account for this unexpected donor by including their genotypes among those of the other candidate donors in Census-seq analysis; this analysis revealed that cells from the CSES15 line had in fact taken over the village (**Figure 3F**). Examination of whole genome sequence data from CSES15 revealed that it harbored an acquired mutation in *MDM4*.

A more challenging analytical problem can arise if contaminating cells come from a donor whose genotypes are unknown. We developed the Contaminating Sample Identifier (CSI) algorithm to detect the presence of contaminating cells from genomically unknown donors (**Figure 3D, E, S2E-G**). CSI utilizes sequence reads that suggest the presence of alleles that are segregating in human populations yet (by chance) absent among the candidate members of a village; CSI determines whether such reads are sufficiently numerous that sequencing error is unlikely to explain them. To evaluate CSI, we asked whether it could identify the presence of contaminating cells in the village described above, but do so in the absence of any *a priori* genetic data for the contaminating CSES15 line. Indeed, CSI analysis suggested the presence of an unexpected donor, at increasing frequency with each passage (**Figure 3E, F**). Follow-up CSI and Roll Call analyses of individual cell lines (from which the village was made) allowed us to identify the tube that had initially become contaminated with CSES15.

Roll Call and CSI can be used to authenticate cellular reagents – for individual cell lines as well as villages – in a wide variety of laboratory contexts.

### Cellular phenotyping and genetic analysis in villages

To analyze how a cellular phenotype of interest varies among donors, we analyze how sorting or selection for that phenotype changes the representation of each donor’s DNA in the cell mixture (**Figure 4**). Although a donor’s individual cells may also vary in phenotype, donor-to-donor variation in the mean and/or variance of such distributions will alter the representation of each donor’s DNA in the resulting villages (**Figure 4A**). For example, to analyze the expression level of a specific protein, we stain about 10^6^-10^7^ cells from a cell village with an antibody to that protein, then sort the cells based on the immunofluorescence signal (**Figure 4A, 4B**). Each donor’s quantitative phenotype is then the ratio of their DNA representation in two different selected villages (**Figure 4C**).

**Figure 4.**
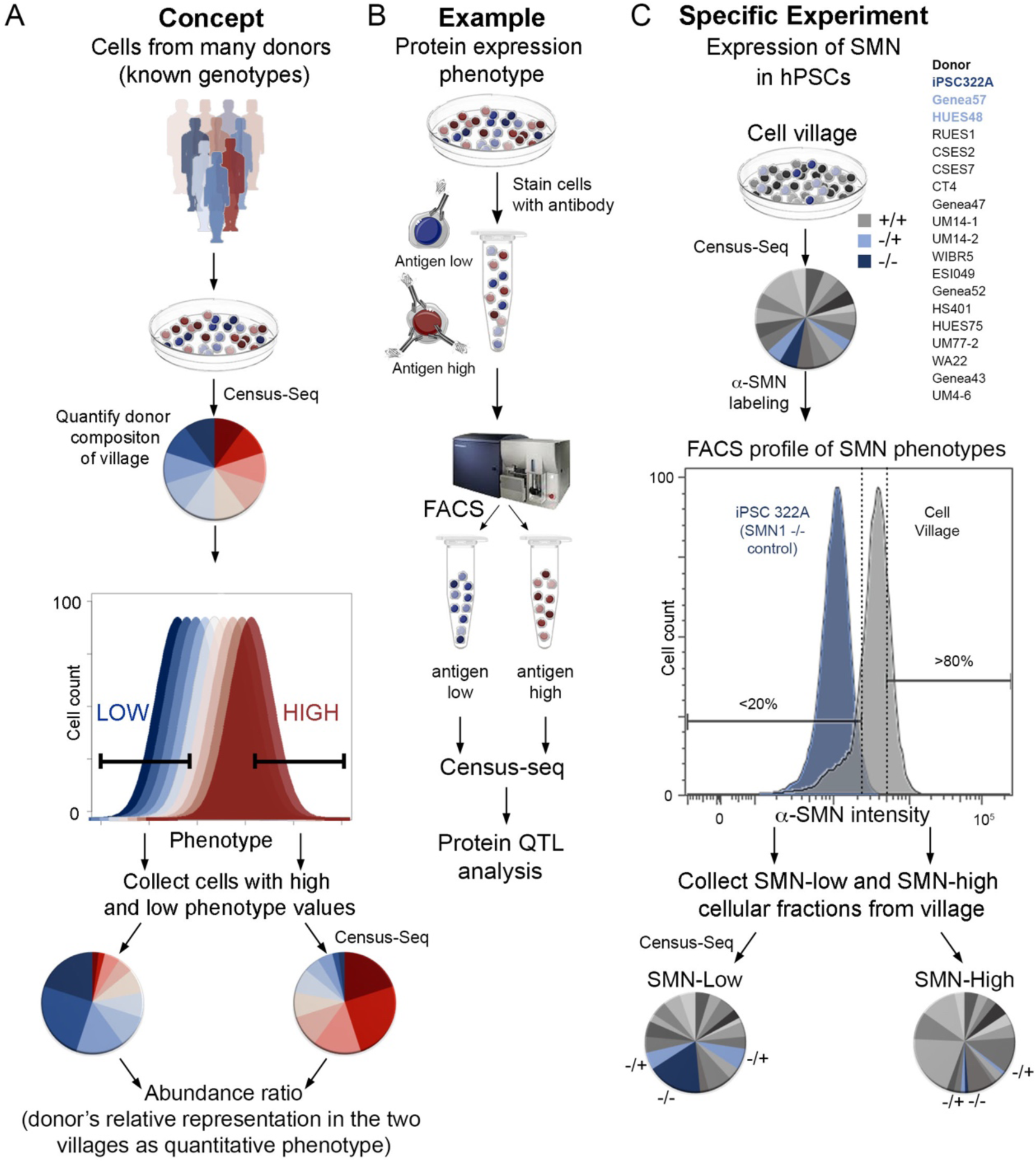
Using cell villages and Census-seq to analyze cellular phenotypes at population scale. A. A cellular phenotype may be affected by both inter-individual biological variation and single-cell variation (in biology or measurement). Sorting or selecting the cells in the village based on this phenotype creates derived villages. If inter-individual biological variation shapes this cellular phenotype, then the derived villages will have different donor compositions. A donor’s change in representation between such derived villages is a quantitative phenotype that can then be analyzed genetically or in relationship to other variables, such as donor age or health status. B. Ascertainment of inter-individual variation in protein expression from cell villages using Census-seq. A cell village is fixed and stained with an antibody to a protein or post-translational modification of interest. The cell village is FACS-sorted for level of immunoreactivity. The donor composition of each cell fraction is analyzed by Census-seq. C. A cell village was analyzed for expression levels of the SMN protein. The pilot village, consisting of PSCs from 19 donors, included cells from a donor with spinal muscular atrophy (SMA, a recessive genetic disorder caused by SMN deficiency) and two carriers of recessive SMA mutations. Comparisons of the FACS-derived SMN-high and SMN-low cell villages by Census-seq indicated that cells from all three donors were more abundant in the SMN-low than the SMN-high fraction. This effect was strongest for cells from the SMA patient, and also detectable in the two carriers.

The ability to analyze DNA recovered from fixed, sorted cell mixtures enables many kinds of analyses. Intracellular as well as cell-surface proteins can be analyzed. Flow cytometry conditions and gating thresholds can be optimized, for example by using mutant cells, strongly perturbed cells, or secondary-antibody-only conditions to define reference distributions of phenotypic values (**Figure 4C, S3**). Aliquots of a cellular village, once prepared and fixed, can be used to analyze many different proteins.

As a model cellular phenotype, we analyzed genetic influences on the Survival of Motor Neuron (SMN) protein, which in humans is encoded by the paralogous *SMN1* and *SMN2* genes on chr5q13 (Lefebvre et al., 1995, Lorson et al., 1999, Monani et al., 1999). SMN deficiency results in widespread splicing defects and causes Spinal Muscular Atrophy (SMA). Though the coding-sequence differences that distinguish *SMN1* from *SMN2* are all synonymous changes in codon usage, *SMN2* lacks a key splicing enhancer, with the result that the majority of *SMN2* mRNAs produce a shorter protein (*SMNdelta7*) whose inability to rescue *SMN1* deficiency has been attributed to protein instability and perhaps to nonsense-mediated decay (Burnett et al., 2009, Hua et al., 2008). SMN deficiency is primarily caused by mutations in *SMN1*, which are under these circumstances not rescued by *SMN2*. An emerging therapeutic strategy for SMN deficiency is to cause the *SMN2* pre-mRNA to splice in an *SMN1*-like manner, producing a protein that can rescue *SMN1* deficiency (Finkel et al., 2016, Groen et al., 2018, Hua et al., 2010, Meyer et al., 2009, Palacino et al., 2015, Ramdas and Servais, 2020).

To first see whether Census-seq could be used to recognize an individual with a strong SMN-protein-expression phenotype, we created a 19-donor PSC village that included iPSCs derived from an SMA patient (iPSC322A). The cell village was fixed, stained with a monoclonal antibody recognizing the SMN protein (produced by both the *SMN1* and *SMN2* genes), and sorted based on anti-SMN immunoreactivity (**Figure 4B, C**). We then separately collected cells whose immunoreactivity was in the upper and lower quintiles relative to the rest of the village (**Figure 4C, Figure S3**). Census-seq comparison of these “SMN-immunoreactivity-high” and “SMN-immunoreactivity-low” villages revealed that, as expected, cells from the SMA patient were greatly over-represented (3.1-fold) in the SMN-low fraction relative to the SMN-high fraction (**Figure 4C**). Interestingly, cells from two additional cell lines were also over-represented in the SMN-low fraction (1.9-fold and 1.4-fold, **Figure 4C**). Examination of WGS data from the 19 cell lines in the village revealed that these two cell lines came from heterozygous carriers of an *SMN1* deletion.

In addition to the SMA patient and two *SMN1* deletion carriers, the other 16 PSC donors also varied in their DNA contribution to the SMN-low and SMN-high cell villages (**Figure 4C**). To see whether such variability was driven by genetic variation, we analyzed this phenomenon in a larger village of 113 iPSC lines obtained from the California Institute of Regenerative Medicine (**Figure 5A**), whose genomes we also analyzed by WGS and make available as part of this work (Lin et al., 2020). Census-seq revealed abundant variation in the SMN-expression phenotype (**Figure 5A**) even among donors whose WGS data indicated that they did not harbor heterozygous or homozygous deletions in *SMN1*. To determine the genetic source of this phenotypic variation, we used the Census-seq measurements to perform a genome-wide association study (GWAS) of the SMN-expression phenotype – which we scored as the log-ratio of each donor’s representation in the SMN-high and SMN-low fractions – focusing on 96 iPSC lines that were sufficiently well-represented in the villages to yield high-precision measurements of their representations (**Figure 5B, 6D, blue symbols; Methods**). This analysis yielded a genome-wide significant (p = 2.91×10^−14^) association to the locus on chromosome 5 containing the *SMN* genes. The phenotype mapped most strongly to common copy-number variation of the *SMN1* and *SMN2* genes. We found most individuals had inherited 2 to 6 such genes (total), which we measured in each donor by applying the Genome STRiP algorithm to the individual donors’ WGS data (**Figure 5C, 7B**) (Handsaker et al., 2011, Handsaker et al., 2015). The strong statistical significance of this relationship – which resulted from both the number of donors in the analysis (96) and the high correlation of gene copy number with the Census-seq SMN phenotype measurements (*r*^*2*^ = 0.59) – meant that the genetic basis of this phenotype could in principle have been mapped in an unbiased genome-wide search.

**Figure 5.**
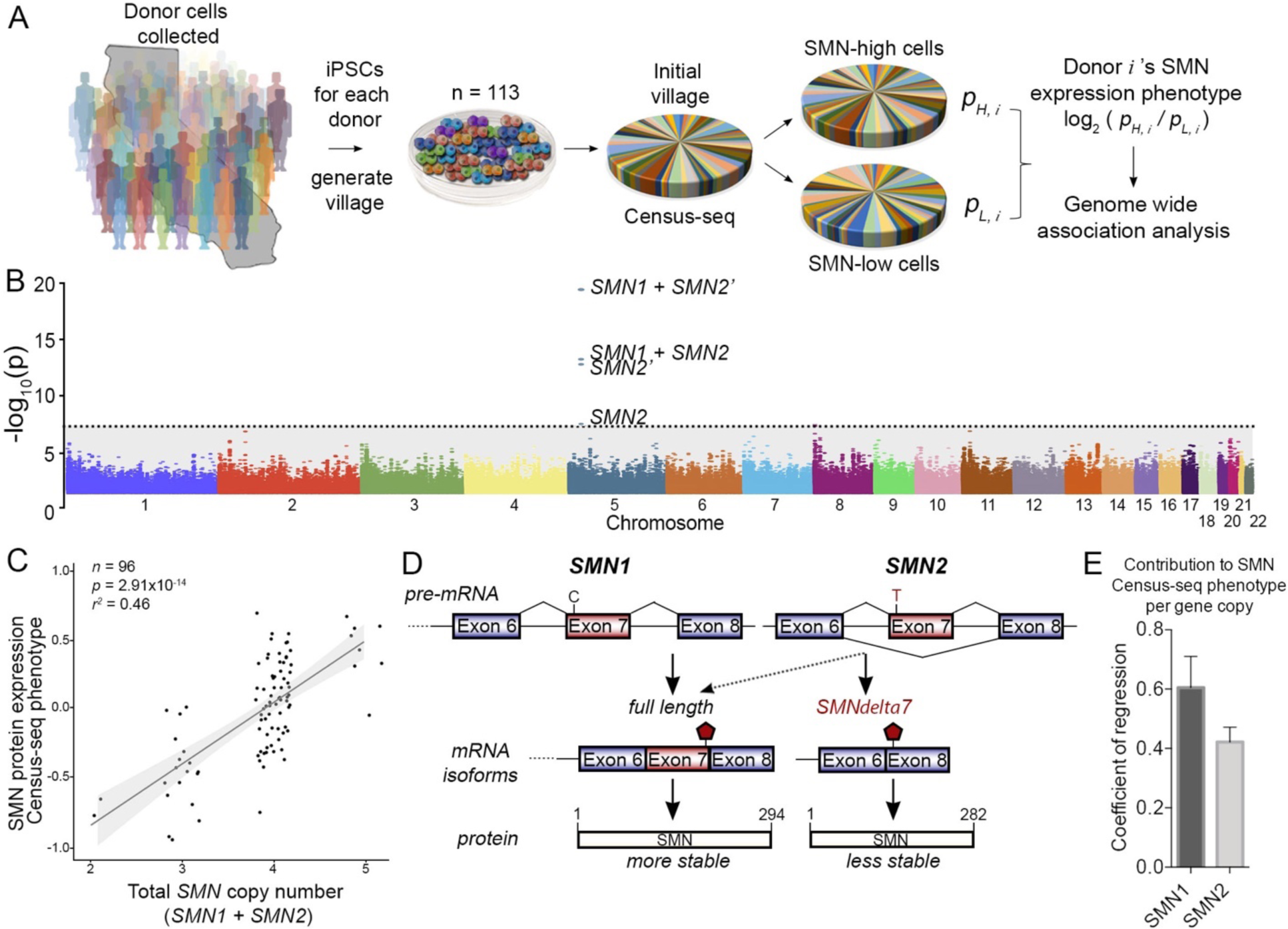
Genetic basis of an SMN protein expression phenotype. A. A village of iPSCs from 113 donors was assembled. The cells in the village were fixed, immunostained for SMN protein, and sorted into SMN-high and SMN-low fractions. Genomic DNA from the two fractions was analyzed by Census-seq. For genetic analysis, each donor’s SMN-expression phenotype was quantified from the relative abundance of his/her genomic DNA (cells) in the SMN-high and SMN-low fractions. B. Manhattan plot showing genome-wide association analysis of this SMN protein-expression phenotype. This analysis revealed genome-wide-significant association at the locus containing the *SMN1* and *SMN2* genes, which exhibit common variation in gene copy number. Genome-wide-significant associations involved *SMN2* gene copy number and (more strongly) the combined copy number of *SMN1* and *SMN2. SMN2’* refers to a calculation of *SMN2* gene copy number that excludes a potential null allele characterized in Figure. 7. C. Correlation of the Census-seq SMN protein-expression phenotype with the summed gene copy number of *SMN1* and *SMN2* genes. D. Though *SMN1* and *SMN2* encode identical amino acid sequences, a sequence variant in a splice enhancer causes many *SMN2* mRNAs to splice in a way that excludes exon 7, resulting in a protein that is less stable and potentially less functional. E. *SMN1* and *SMN2* contribute unequally to the Census-seq SMN protein-expression phenotype. Bars indicate coefficients of a linear regression of the SMN protein-expression phenotype against *SMN1* and *SMN2* gene copy numbers. Error bars indicate standard error.

Each copy of *SMN2* can only partially rescue loss of an *SMN1* allele – despite being expressed in the same tissues and cell types – a failure that could in principle be due to differences in the stability or activity of the proteins generated by *SMN1* and *SMN2* (**Figure 5D**). To separately quantify the contributions of *SMN1* and *SMN2* to SMN protein abundance, we used the fact that *SMN1* and *SMN2* each exhibit common variation in copy number; we inferred each donor’s gene copy number for *SMN1* and *SMN2* by utilizing paralogous sequence variants that distinguish between the genes (**Methods**). Linear regression of the donors’ Census-seq SMN-expression phenotypes against their *SMN1* and *SMN2* gene copy numbers revealed that both *SMN1* and *SMN2* copy number contributed positively to SMN protein expression, with *SMN1* making a greater contribution (**Figure 5E**). This result confirms that *SMN1* generates somewhat more or longer-enduring SMN protein than does *SMN2*. However, the modest difference between the per-copy effects of *SMN1* and *SMN2* (**Figure 5E**) suggests that protein instability is not on its own a sufficient explanation for the inability of *SMN2* to rescue *SMN1* deficiency and is consistent with the hypothesis that *SMNdelta7* also has reduced activity.

### Pharmacogenetics in cell villages

An important therapeutic approach is to coax *SMN2* to splice in an *SMN1*-like manner, generating a more stable and/or more effective protein. The clinical candidate LMI070 (Branaplam) was identified in a screen for modulators of *SMN2* splicing (Cheung et al., 2018, Singh and Singh, 2018). LMI070 appears to interact with the splicing enhancer in exon 7 and to increase levels of SMN protein in cells in a concentration-dependent manner (**Figure 6A**). The efficacy of LMI070 for treating SMA is currently being evaluated in phase I/II clinical trials in multiple countries (<u>NCT02268552</u>).

**Figure 6.**
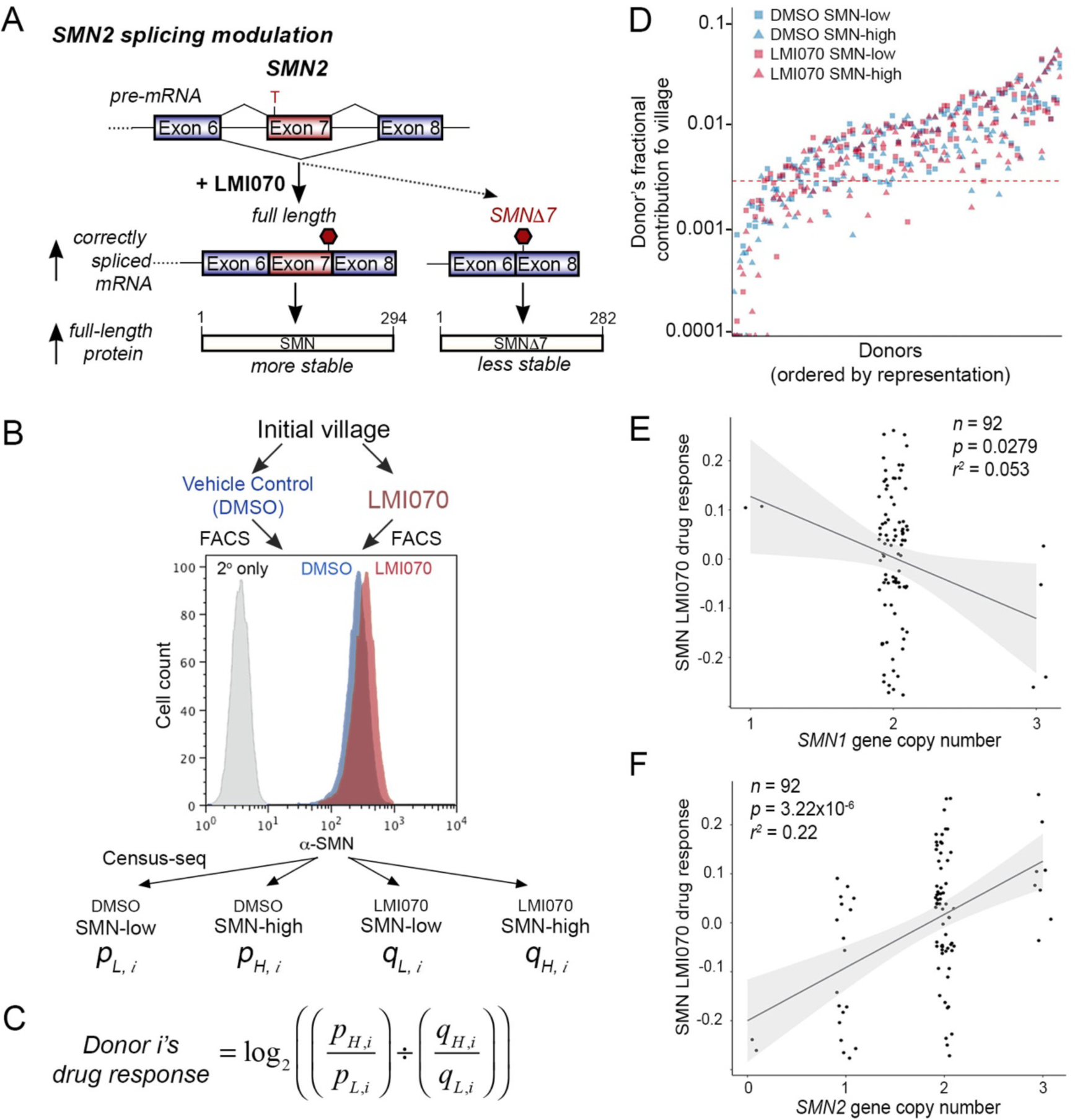
Pharmacogenetic analysis of response to SMN therapy. A. An emerging therapeutic approach for SMA is to cause *SMN2* to splice in an *SMN1*-like manner. The drug LMI070 was developed to do this. B. An iPSC village of 113 donors was split into two villages, which were then treated with either LMI070 or a vehicle control (DMSO). Both the LMI070-treated and the vehicle-treated villages were then fixed, immunostained and sorted into SMN-high and SMN-low fractions. C. For genetic analysis, a donor’s LMI070-response phenotype was calculated from her/his relative cellular contributions to these four fractions. D. Estimated contribution of each of the 113 donors to each of the four derived villages. E. Correlation of the Census-seq LMI070-response phenotype with *SMN1* gene copy number. F. Correlation of the Census-seq LMI070-response phenotype with *SMN2* gene copy number.

As humans often vary in drug responses, a key need in biomedical research is to be able to anticipate individuals’ response to drugs and predict who might have an optimal or non-optimal response. We sought to understand whether cell villages could be used to identify variation in drug response and uncover genetic contributions to drug response. We first found concentrations of LMI070 that could cause changes in SMN protein expression (full-length and *SMNdelta7*) across three cell lines with varying gene copy number of *SMN1* and *SMN2* (**Figure S4**). Villages of iPSCs were then exposed to either LMI070 (0.1uM) or a vehicle control (DMSO) for 24 hours (**Figure 6B**) before being sorted into SMN-high and SMN-low fractions. We measured each donor’s relative-drug-response phenotype by calculating how LMI070 treatment changed the distribution of that donor’s DNA into the SMN-high and SMN-low cell fractions, relative to the vehicle control (DMSO) (**Figure 6C, Methods**); this drug-response phenotype variable was calculated from the results of four Census-seq analyses (**Figure 6B, C**).

Cells’ LMI070-response phenotype correlated strongly with gene copy number of *SMN2* (*p* = 3.22×10^−6^) but not *SMN1* (*p* > 0.01), consistent with the hypothesis that LMI070 affects SMN protein levels by acting specifically upon *SMN2* (**Figure 6E, F**). These results replicated in a distinct village of hESCs (**Figure S5**). These results contrasted strongly with the baseline SMN-expression phenotype, which was affected more strongly by *SMN1* than *SMN2* variation (**Figure 5E**).

Because *SMN1* deficiency strongly affects neurons, we also characterized the LMI070 response phenotype in villages of neural cells. A village of neural cells (from 50 donors) was generated using a lentiviral-based delivery system to induce the expression of *Ngn2* (**Figure S6**) and treated with LMI070 for 24hr before Census-seq analysis. Both the SMN-expression phenotype and LMI070 drug-response phenotype associated with *SMN2* copy number (**Figure S6C, D**; *p* = 2.36×10^−4^, *p* = 1.15 × 10^−3^), replicating the results from the iPSC village.

Despite the large apparent effect of *SMN2* gene copy number on response to LMI070, we found that cells from donors with the same number of *SMN2* gene copies often exhibited quite different response to LMI070 (**Figure 6F)**, potentially reflecting additional genetic effects. To better ascertain the full spectrum of DNA variation at the *SMN1/SMN2* locus, we analyzed WGS data from 767 individuals (**Methods**). We found that many donors carried an apparent deletion of *SMN2* exons 7 and 8, including the LMI070 binding site (**Figure 7A, B, C**; we refer to this allele as “*SMNdel*” below). Although the deletion was present in the genomes of 10% of the sampled individuals with European ancestry, it has only recently been described (Vijzelaar et al., 2019).

**Figure 7.**
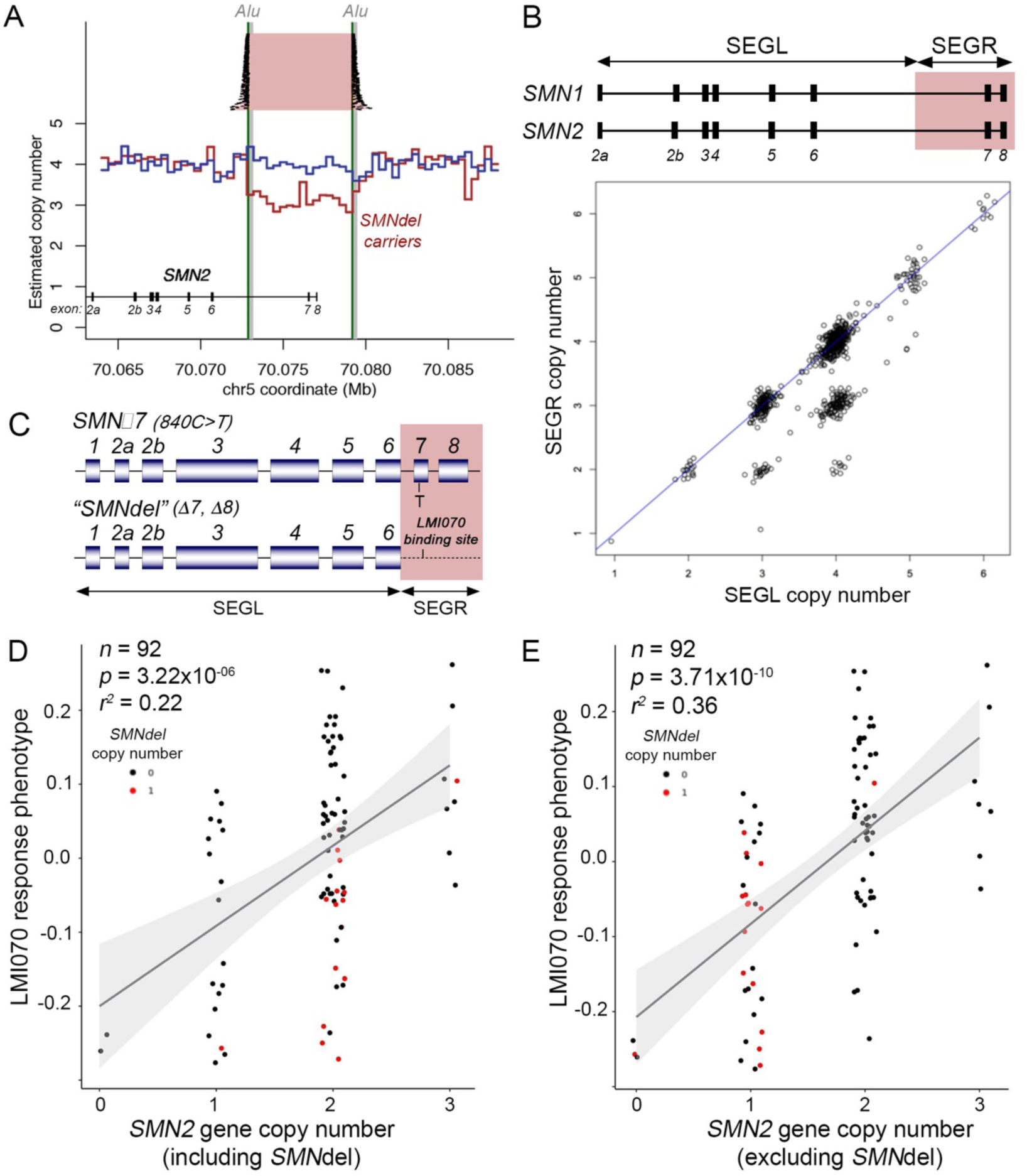
Pharmacogenetic analysis of response to SMN therapy. A. In analysis of whole genome sequence generated from genomic DNA from the individual iPSC donors, 22 of the 113 donors were found to have sequence reads or mate-paired reads that appeared to jump over a 6-kb genomic segment (flanked by two Alu repeats) containing two exons of the *SMN2* gene. Those same individuals tended to have reduced copy number of the 6-kb segment (red) relative to the other individuals (blue), as estimated from read depth of coverage across the genomic locus. B. Across 767 genomes analyzed by whole genome sequencing, copy number of this 6-kb segment (SEGR) was in many individuals less than copy number of the rest of the *SMN1/SMN2* gene (SEGL). (Note that the >99% sequence identity between *SMN1* and *SMN2* requires that the paralogous sequences from these two genes be counted together for this analysis.) This population-level pattern confirms that SEGR is affected by a cryptic, common deletion allele. C. This cryptic, common deletion allele (“*SMNdel*”) removes two exons, including the exon that encodes the putative binding site for LMI070. D. Cells from individuals with the *SMNdel* allele in their genomes tend to have a smaller response to LMI070 (as measured by the Census-seq LMI070-response phenotype) relative to cells from other individuals with the same *SMN2* gene copy number. Red points: *SMNdel* carriers. Black points: Other iPSC donors. E. The Census-seq LMI070-response measurements more strongly fit a model in which *SMNdel* is treated as a null allele of *SMN2*.

Re-analysis of the Census-seq data to account for the *SMNdel* allele showed that carriers of *SMNdel* were indeed the donors whose cells exhibited a weak response of SMN expression to LMI070 treatment (**Figure 7D, red**). The Census-seq LMI070-response phenotype correlated much more strongly with a measure of “intact” *SMN2* gene copy number that we obtained by treating SMNdel as a null allele (*r*^2^ = 0.36, *p* = 3.71^-10^), suggesting that SMNdel carriers under-respond to LMI070 (**Figure 7E**). In fact, we found evidence that *SMNdel* is either not translated, or encodes a much less stable protein than *SMN2* does: excluding *SMNdel* copies from genetic measurements of *SMN2* gene copy number greatly strengthened the association with the SMN baseline protein-expression phenotype (**Figure 5B**). Simultaneous linear regression of the Census-seq SMN-expression and LMI070-response phenotypes against gene copy numbers of *SMN1, SMN2* (excluding *SMNdel*), and *SMNdel* **(Figure S7**) indicated the *SMNdel* allele, in addition to encoding an LMI070-unresponsive RNA, also encodes a much less stable isoform of the SMN protein than canonical *SMN2* gene copies do.

## Discussion

Genetic variation shapes almost all human phenotypes, creating a profound opportunity for biological discovery and translational biology if science can begin to reveal how human alleles – individually, and in concert – shape the life of cells. To help realize this scientific possibility, we developed “village-in-a-dish” experimental systems, in which cells from scores of donors are grown, perturbed and phenotyped in a single reaction chamber; these systems enable population-genetic approaches to cell-biological questions. The analysis of these systems is enabled by three computational methods – Census-seq (**Figure 1**), Roll Call (**Figure 3**), and CSI (**Figure 3**) – which reveal and learn from the donor composition of cell and DNA mixtures. The practical execution of such systems is further enabled by ways to create and maintain high-quality cell villages.

Human genomes teem with functional variation. Here, even at a single locus, Census-seq analyses revealed effects of at least three kinds of genetic variation. These included (i) *SMN1* gene copy number, which affected SMN protein levels at baseline but not responsiveness to LMI070 therapy; (ii) *SMN2* gene copy number, which affected SMN protein levels at baseline and also response to LMI070; and (iii) a cryptic *SMN2* allele (*SMNdel*), common but not routinely screened for in clinical diagnostics, which compromised SMN protein levels and abrogated LMI070 response (**Figure S7**).

Since our goal in describing these systems is to make them as facile and adaptable as possible, we here discuss practical lessons for creating and maintaining villages and designing larger research strategies to utilize them.

Variation in the relative growth rates of cells from different donors presents both an interesting area of scientific inquiry and a day-to-day practical challenge for cell villages. By making it easy and inexpensive to collect population information at every cell passage, Census-seq makes it possible to measure proliferation phenotypes at a population scale, to screen hundreds of lines for growth phenotypes and growth-promoting mutations, and to quantify the effects of such mutations (**Figure 2**). When using Census-seq to address other biological questions, however, a natural emphasis will be on containing such population dynamics. We describe **(Methods**) practical approaches for doing this, including sequencing to identify acquired mutations, designing small villages (“heats”) to identify empirically hyper-proliferative cell lines (**Figure 2**), and – for experiments on post-mitotic or differentiated cell types – quickly moving PSC villages into post-mitotic states (**Figure 2**).

Though cell villages and populations enable primary genetic discoveries in Census-seq, it is useful in many contexts to analyze individual control cell lines alongside villages, and to seed villages with one or more lines already known to harbor strong effects. The choice of sorting or selection criteria in Census-seq experiments can be enhanced by comparison to one or more individual cell lines expected to have a strong phenotype. For example, flow-cytometric analysis of cells from an SMA patient informed the flow-cytometric gating strategy that we then applied to the village for population-scale genetic analyses of SMN protein expression (**Figures 4, 5, S3**). Positive- and negative-control lines can also be derived by methods including gene editing CRISPR-i, CRISPR-a, and pharmacological perturbation of individual lines (Gasperini et al., 2019, Larson et al., 2013, Hsu et al., 2014). The further inclusion of such lines in villages enables the effects of population-genetic variation to be compared to the effects of strong laboratory perturbations. The ability to detect expected, positive-control effects also serves as a useful gate establishing that an experiment has been executed successfully. The inclusion of such positive controls may also allow the lack of variation across scores of other cell lines to be a meaningful statement about biological constraint on a cellular phenotype.

Our finding that cells from carriers of the *SMNdel* allele responded less strongly to LMI070 raises an interesting issue that will need to be addressed for many drug candidates that target RNA metabolism. Polymorphisms that influence RNA structure and regulation may be much more abundant than polymorphisms that influence the peptide sequence of the encoded protein, the traditional drug target. Census-seq could be used to identify, prior to clinical studies, individuals and genotypes who would have optimal and/or unexpected responses to such therapeutic candidates.

Additional challenges may arise in other experimental contexts. Mitotic cells present challenges in population dynamics (**Figure 2**) that were managed by the techniques we describe here – including genetic screening (for acquired mutations) and empirical growth-rate measurement (in cellular “heats”). Similarly, variability in pluripotency after reprogramming or exposure to various environmental conditions across cells’ lifetime may influence their differentiation potential (Cahan and Daley, 2013, Nishizawa et al., 2016). The refraction of cells down a particular pathway may even be shaped by genetic variation, a potential area for Census-seq analyses. Finally, a central challenge to the maintenance and expansion of cellular resource banks that are required to facilitate population-scale experiments has been detecting cell line contamination and validating of donor identity (Liang and Zhang, 2013, Rouhani et al., 2014).

These issues have historically been of major concern and have real implications on experimental reproducibility (Neimark, 2015, Nelson-Rees et al., 1981). The methods described here (including Census-seq, Roll Call and CSI) create a greatly expanded and inexpensive tool kit for validating the identity and purity of cellular resources.

Village-in-a-dish systems make a tradeoff between detection of cell-autonomous and cell-nonautonomous genetic effects. By analyzing cells from all donors together, Census-seq normalizes most cell-non-autonomous effects across donor genotypes. Most of these cell-nonautonomous effects are common sources of experimental noise – for example, effects of cell density and its downstream effects on metabolite concentrations, cellular waste-product concentrations, cell-cell contacts, and (for neurons) synaptic stimulation and activity. The normalization of such effects in Census-seq makes cell-autonomous effects more visible above experimental noise; this was evidenced by the strong explanatory power of common alleles in almost all of our experiments (**Figures 5-7**) and the ability of these experiments to establish genotype-phenotype relationships at genome-wide statistical significance. In many contexts, though, cell-nonautonomous effects may be of real interest, so it is important to think about how to design Census-seq experiments to ascertain them. Census-seq could be readily used to map cells’ responses to a non-cell-autonomous stimulus such as a ligand. However, different kinds of analysis would be needed to map genetic effects on the magnitude of the stimulus itself; this might be possible in Census-seq if the ligand’s synthesis can be made the subject of a Census-seq analysis.

A key question for population-scale cell biology and systems-biological modeling involves the ability to discover and quantify modest, quantitative effects. The experiments described here ascertained strong, rare, Mendelian effects (as in cells from an SMA patient) and more modest, common effects, such as effects of common variation in gene copy number on protein expression and therapeutic response. Key to the ability to discover and characterize modest, quantitative effects is the ability to equalize technical factors across donors, reducing experimental noise and thus increasing the detectability of genetic signals relative to noise. The specific parameters chosen for selections and screens are also important and are worth careful thought and optimization. For example, the flow-cytometric gates used to select for derived villages (**Figure S3**) can be critical for measuring genetic effects with high sensitivity. Since Census-seq molecular and computational analyses are inexpensive (sequencing expenses are about $100 per village), we encourage researchers to experiment with a variety of parameters and observe the impact of these variables on measurement of known, positive-control effects.

Complex phenotypes with multi-locus inheritance will present interesting challenges to the design of Census-seq experiments, often requiring analysis in still-larger numbers of donors. We anticipate that the optimal approach to this will involve analyzing multiple villages each of 60-120 donors, with modest overlapping membership to inform meta-analysis. In our experience, most of the returns of Census-seq in scalability and well-controlled measurement are already realized at the scale of 40-120 donors – a level at which village assembly can also be performed by a single scientist in a single session. Villages of 40-120 donors might provide a natural unit of larger meta-analyses that seek to find genetic effects that are rare or modest in magnitude.

In selecting more complex phenotypes for study, a promising next direction may be in analyzing oligogenic phenotypes that might be affected by variation at a few loci, perhaps focusing on those pathways or molecular complexes in which genome-wide association studies (by SNP arrays or exome sequencing) have already identified risk variants or haplotypes in several different genes. Such constellations of (common and rare) genetic effects may suggest a natural integration point for cellular-genetic analysis. In schizophrenia, for example, many different genetic associations involve subunits of the L-type calcium channels, whose expression, membrane localization or even physiology could be made the subject of cellular screens and selection (Lam et al., 2019, Schizophrenia Working Group of the Psychiatric Genomics, 2014).

Polygenic phenotypes will ultimately require the largest samples and present the greatest challenge – and perhaps also the greatest reward, since such polygenetic architectures have been extremely challenging to dissect by traditional biological methods and are hypothesized to involve indirect effects through complex cellular networks (Boyle et al., 2017). An exciting possibility is that, even for polygenic illnesses, it may be possible to find cellular phenotypes that are potential convergence points of genetic effects whose functional connection was not previously appreciated. Such cellular phenotypes might associate in villages with donors’ polygenic risk scores, which for many complex phenotypes identify individuals with risk equivalent to that of well-known monogenic mutations (Khera et al., 2018). Villages can in principle be designed from individuals in the two “tails” of a polygenic risk-score distribution – i.e. who have extremely high or low polygenic risk scores. It is challenging to predict how many donors will be required for such analyses, as any prediction presumes an answer to a central unknown question – the extent to which different genetic effects on a disease phenotype will converge upon a few key cellular processes. We hope population-scale experimental systems present an empirical, data-driven path toward answering these and many other questions.

## Acknowledgements

Support for this work has been provided by the Broad Institute’s Stanley Center for Psychiatric Research and, since 2017, by a “Convergent Neuroscience” grant from the National Institutes of Mental Health (U01 MH115727, to SM and KE). This study used induced pluripotent cell lines generated by the California Institute for Regenerative Medicine (CIRM) and distributed by Cellular Dynamics International (CDI). We thank CIRM and CDI for sharing genotype and sample metadata for these iPSCs with us.

## Author Contributions

Conceptualization and design: JMM, JN, SG, KE, SM; Experimentation: JMM, CM, KR, HdR; iPSC line expansion, QC and sequencing: MT and DH, supervised by RN. Data analysis: JN, REH, SG, SM, DM; Supervision: KE, SM; Project administration: AN

## Supplementary Figures

**Figure S1.**
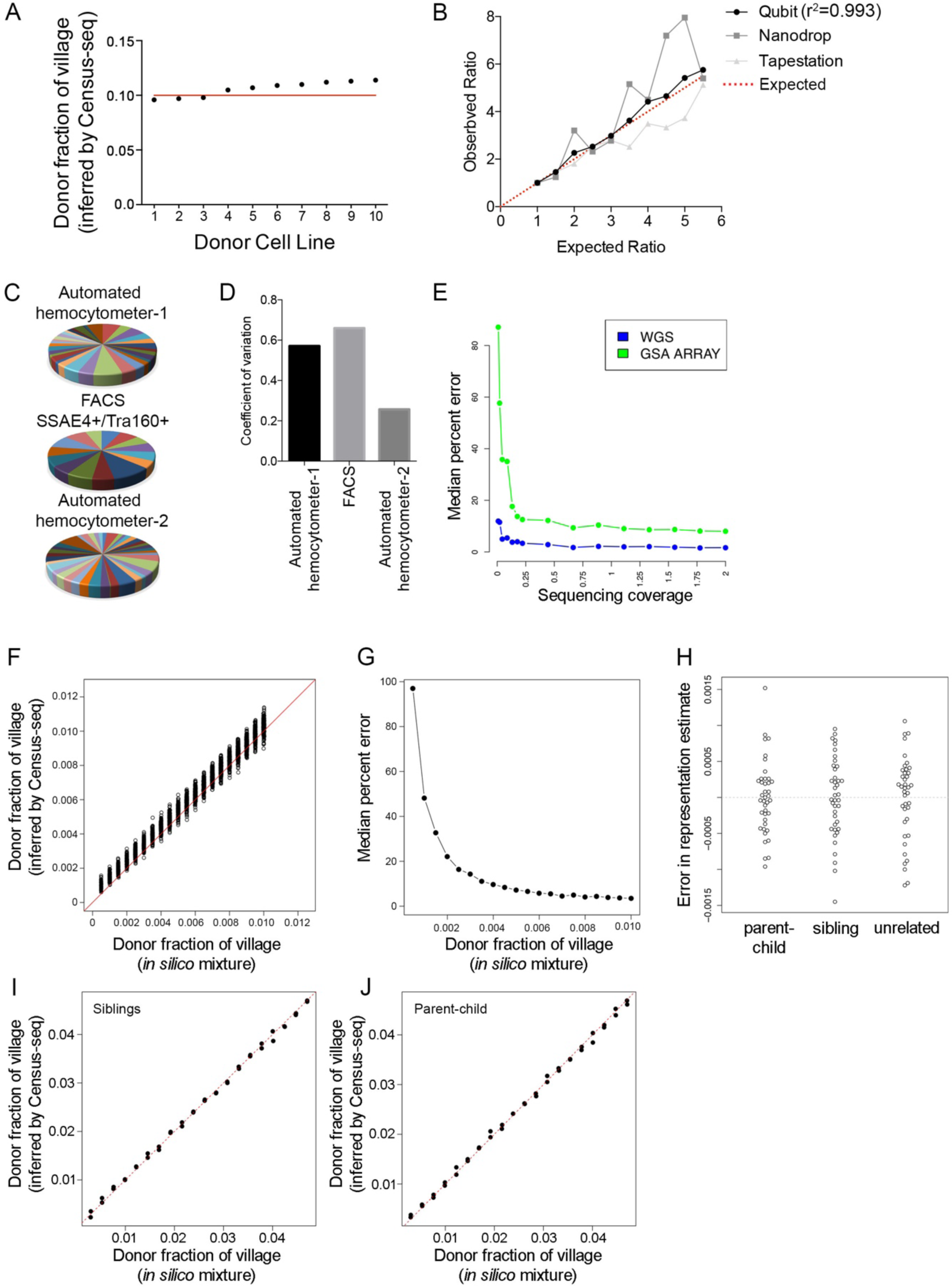
Construction and quantification of cell villages. A. Equal quantities of genomic DNA from 10 donors was mixed, sequenced and analyzed by Census-seq. The plot compares the aliquoted donor-contribution proportions of the DNA mixture to the Census-seq estimates from the sequencing data. B. Genomic DNA from six donors was mixed in an arithmetic progression of quantities as estimated by three DNA-quantitation technologies (Qubit, Nanodrop, Tapestation). Donor contribution to the resulting DNA mixtures was then estimated by Census-seq. Census-seq estimates of donor-specific contributions to the mixture agreed most strongly with input estimates from the Qubit (*r*^2^ = 0.993). C. Cell lines were mixed using three different methods of cell quantification, with the goal of making equimolar mixtures; pie charts show the resulting donor composition of each mixture as estimated by Census-seq. The three cell-counting methods were: a hemocytometer (Countess, for live/dead estimation; flow cytometry with selection for SSEA4 and Tra-1-60 positive and pluripotent) cells; and an automated cell counter (Scepter) based on cell size. D. Using the data from (D), the coefficient of variation (standard deviation divided by the mean) of the donor contributions was estimated for each of the methods of cell quantification. E. Precision of the Census-seq algorithm as a function of (i) the type of *a priori* genome-variation data available for each donor (line colors) and (ii) the depth to which the village genomic DNA is sequenced (y-axis). A key relationship is that deeper *a priori* genetic analysis (e.g. WGS, blue curve) of the individual donors’ genomes makes it possible to infer the donor composition of mixtures even from sequence data that are relatively low-coverage. Analysis is based on a WGS data mixed from 40 donors. F. Results of *in silico* data-mixing experiments to estimate the bias and variance of Census-seq inferences for donors who have contributed small proportions (0.05% to 1%) to a mixture. WGS data from 40 unrelated donors was mixed *in silico*, with 30 donors at an arithmetic series of representations from 0.05% to 1.00% in 0.05% increments, and 10 donors (for whom data not shown) at higher representations, such that the 40 donors’ representations summed to 1. This was repeated for 10 simulations, in each of which donors were permuted between the low and high representation groups at each iteration to generate a total of 300 observations of each bin. G. Bias of Census-Seq estimates (as a fraction of the estimate) for donors who have contributed small fractions (< 1%) of a cell or DNA mixture. Using the data shown in S1F, the median absolute error in representation was calculated as exp(median (abs(log(donor % representation in silico / donor % representation inferred by Census-Seq)))). Bias in donor representation was substantially greater for donors contributing <0.3% to a mixture. Bias was <15% at a representation of 0.3%, and <10% at a representation of 0.4%. We believe that this bias arises from PCR and sequencing errors, which result in reads that (as a group) tend to bias upward the estimated representation of very-low-contribution donors. For this reason, we exclude from some genetic analyses those donors with contributions of <0.3% to a mixture. H. To assess the potential effect of having genetically related donors in a mixture, three *in silico* mixtures of 40 donors were constructed for: 40 unrelated individuals; 20 parent/child pairs; and 20 sibling pairs. In each analysis, WGS data from these donors was mixed uniformly at a representation of 0.025; the data mixture was then analyzed by Census-seq. Error in representation was calculated as the difference between the known and Census-seq-inferred donor-contribution estimates. 95% of inference were within an absolute error of 0.001. The median absolute error in representation estimates were similar: unrelated 3.4×10^−4^, sibling= 4.1×10^−4^, parent-child= 2.6×10^−4^. I. To evaluate the robustness of Census-seq inference to the inclusion of genetically related individuals in a village, *in silico* data mixing was used to simulate a village of 20 sibling pairs, with the same distribution of representations as in Fig. 1E. J. To evaluate the robustness of Census-seq inference to the inclusion of genetically related individuals in a village, *in silico* data mixing was used to simulate a village of 20 parent-child pairs, with the same distribution of representations as in Fig. 1E.

**Figure S2.**
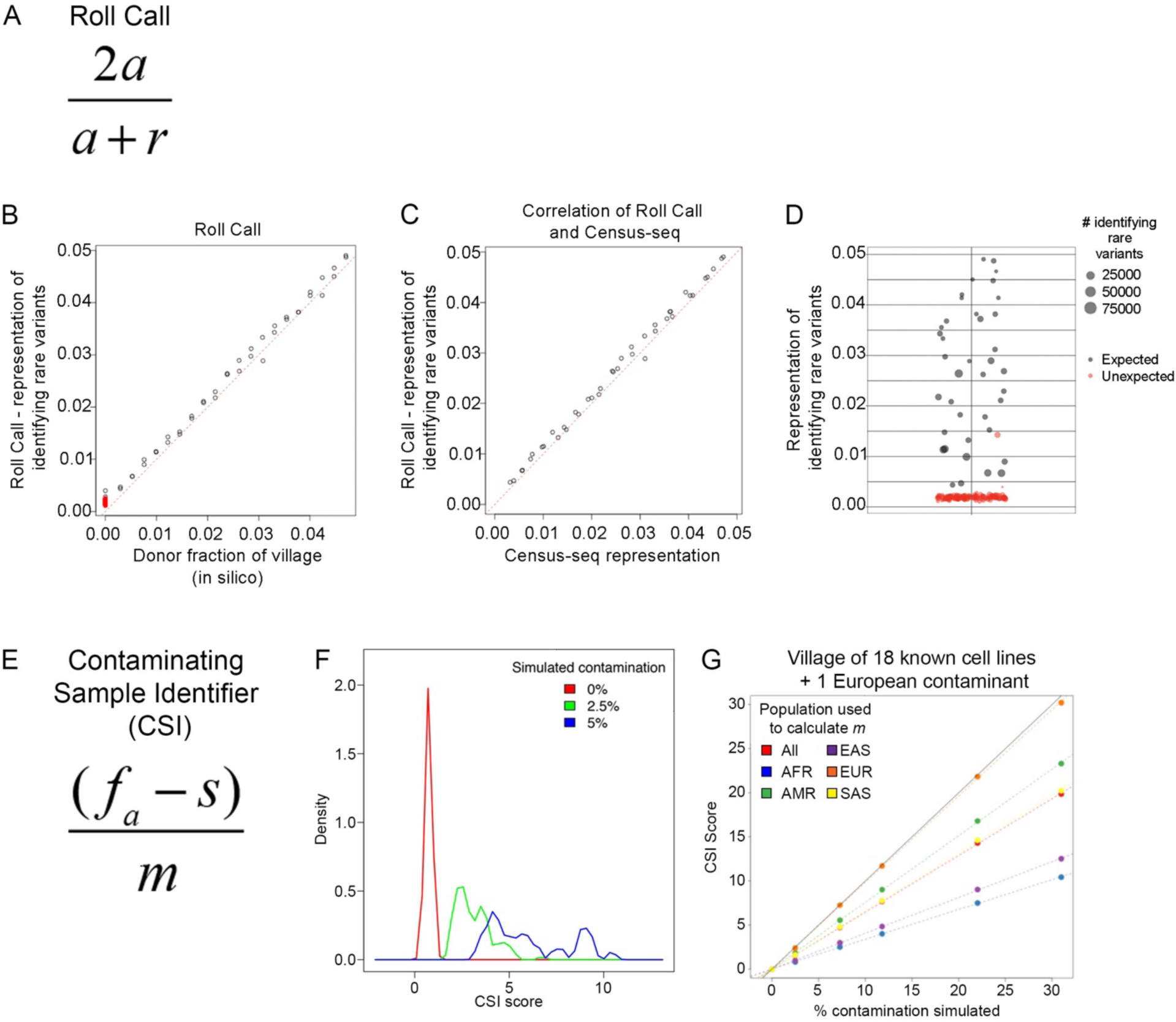
Algorithms and validation analyses for the Roll Call and CSI tools for authenticating cell villages. A. The Roll Call algorithm leverages Identifying Rare Variants (IRVs) – rare alleles that are present in the genome of only one of the candidate donors – to determine which candidate donors have contributed cells/DNA to a mixture. (Census-seq offers more-accurate quantification of donors’ quantitative contributions than Roll Call does, but only when starting with a complete and accurate list of donors.) The formula shows the calculation of the Roll Call score (for an individual donor, using that donor’s IRVs), with which the algorithm evaluates whether a donor has contributed to a DNA mixture. In this formula, *a* and *r* refer to the numbers of observations of the reference and alternate alleles at that donor’s IRVs in the village sequencing data. B. DNA sequence data from 40 unrelated donors was mixed at known proportions. Roll Call scores for these donors (x-axis) are compared to the proportions in which DNA data have been mixed (y-axis). An additional 147 donors (shown in red), whose DNA sequence data was not included in the data mixture, received scores < 0.005 [0.002-0.004]. C. Analysis by Roll Call and Census-seq of a synthetic mixture of WGS data from 24 unrelated individuals, mixed at an arithmetic of contributions across donors. Donors that are known to be in the mixture are observed at close to their correct mixtures. Donors not in the mixture have scores < 0.005 [0.003-0.004]. D. Roll Call analysis of an example cell village confirmed the presence of 45 expected donors (grey circles); confirmed the absence of all but one of the unexpected donors absent (red circles); and flagged the presence of a donor not intended to have been included in that village (red circle at *y*=0.014); follow-up analysis confirmed contamination by cells from this unexpected donor. E. The CSI (Contaminating Sample Identifier) algorithm calculates an intrusion score from sites that are monomorphic among the expected donors (but variable in the wider population from which the donors are sampled) – i.e. from alleles that should not have arisen from the expected donors’ genomes and must therefore represent contaminating cells or sequencing errors. CSI utilizes genomic sites known to vary in the wider population (from which the donors are sampled) that happen to be monomorphic among the candidate donors. *f*_*a*_ is the fraction of allelically informative reads (at these sites) that contain the alternative allele; *s* is the sequencing error rate; *m* is the mean minor allele frequency of these alleles. F. Distributions of CSI scores for *in silico* villages created by WGS data mixing to have 0%, 2.5%, or 5% contamination from an “unknown” donor to whose genetic data the CSI analysis was blinded. The *in silico* mixing experiments each involved a mixture of 39 known, unrelated donors at varying concentrations and (in the 2.5% and 5% cases) an additional randomly selected unrelated donor, to whose genetic data the CSI analysis was blinded. 100 simulated villages per contamination level were analyzed. The 100 null (uncontaminated) simulations yielded CSI scores of 0.0074 +/- 0.0011; CSI scores in all 100 of the 2.5% contamination simulations exceeded any result from this null distribution. G. Results of CSI analyses of synthetic mixtures of WGS data from 18 known donors, plus an additional unexpected donor for whom WGS data was spiked in (to the mixture) at several proportions (“% contamination simulated”, x-axis). Note that because the CSI formula utilizes the mean minor allele frequency of “unexpected” alleles in the sampled population, the CSI intrusion score estimates %contamination correctly only if the intruding cell line is indeed from the population used to estimate this – in this case, 1000 Genomes European-ancestry sample (EUR, orange), since the simulated contaminating line was of European ancestry. When other populations are used, or when the unexpected donor is related to an expected donor (not shown), the intrusion score mis-estimates %contamination. Since the identity of a genomically unknown contaminating line is by definition not knowable *a priori*, the intrusion score is primarily intended to be used as a diagnostic for contamination rather than as a precise measurement of %contamination; however, its change within a village (e.g. over cell passages) in principle reflects change in the proportion of contaminating cells within the village.

**Figure S3.**
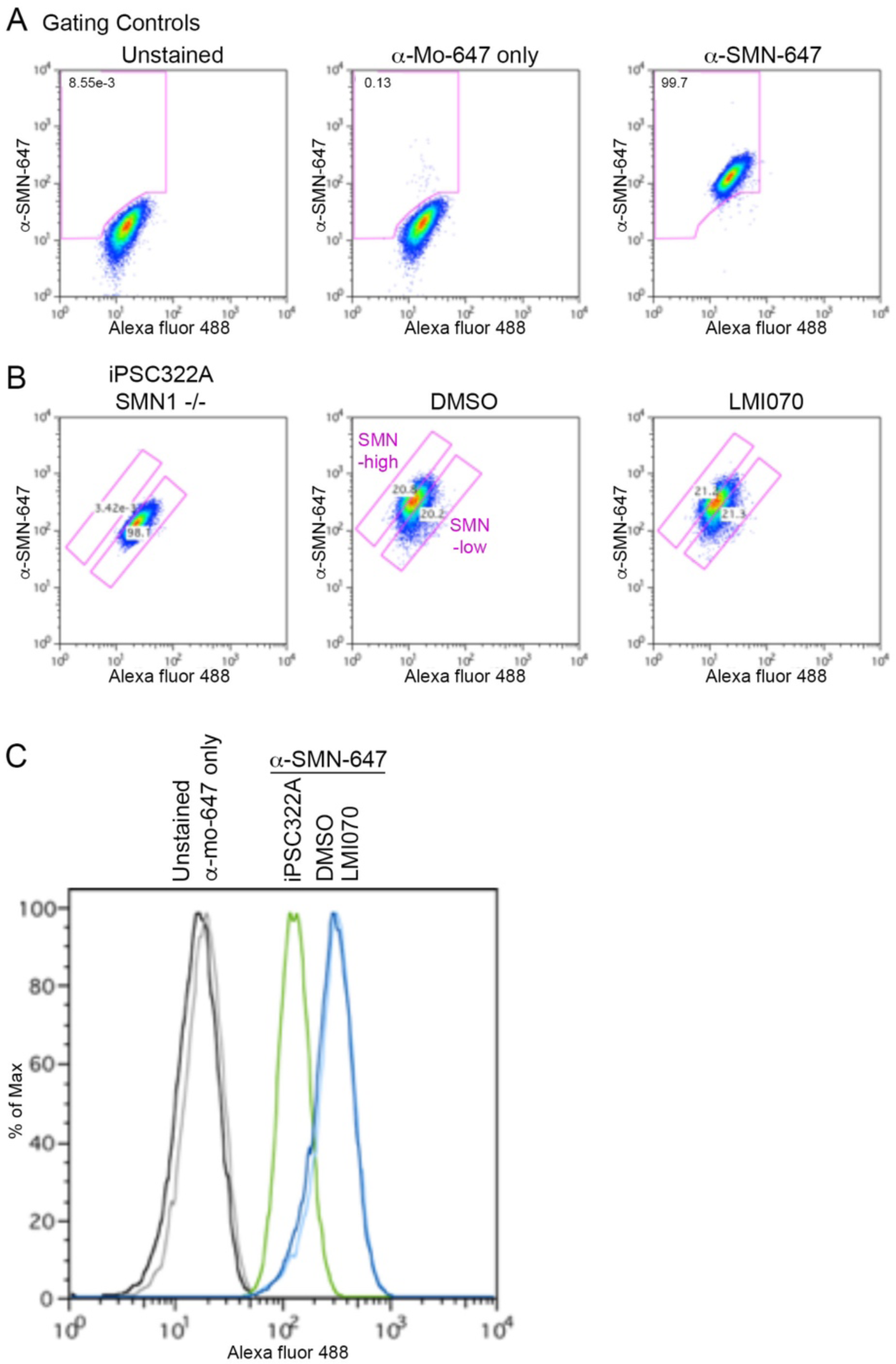
Use of flow cytometry to enrich cell villages for SMN-high and SMN-low cells. A. Cells from a village were dissociated, fixed and immunostained with a monoclonal antibody directed against the protein encoded by *SMN1* and *SMN2*. Gating controls were set using unstained cells (left panel) and cells stained with an anti-mouse Alexa fluor-647 conjugated secondary antibody alone (middle panel). 99.7% of cells stained with the anti-SMN antibody were captured using these gates (right panel). The Alexa fluor-488 channel served as an internal control for autofluorescence. B. Gates were established to capture the top 20% of SMN-stained cells (SMN-high) and the bottom 20% (SMN-low) of cells based on SMN immunoreactivity. Cells from the iPSC322A patient donor line were also analyzed to characterize antibody staining and inform gating (left panel); >98% of cells from the patient cell line. Then, cell fractions were collected from villages treated with a DMSO vehicle control (middle panel) or LMI070 (right panel). C. Histogram representation of the anti-SMN staining data from panels S4A and S4B, to facilitate comparison of these distributions across experiments and conditions.

**Figure S4.**
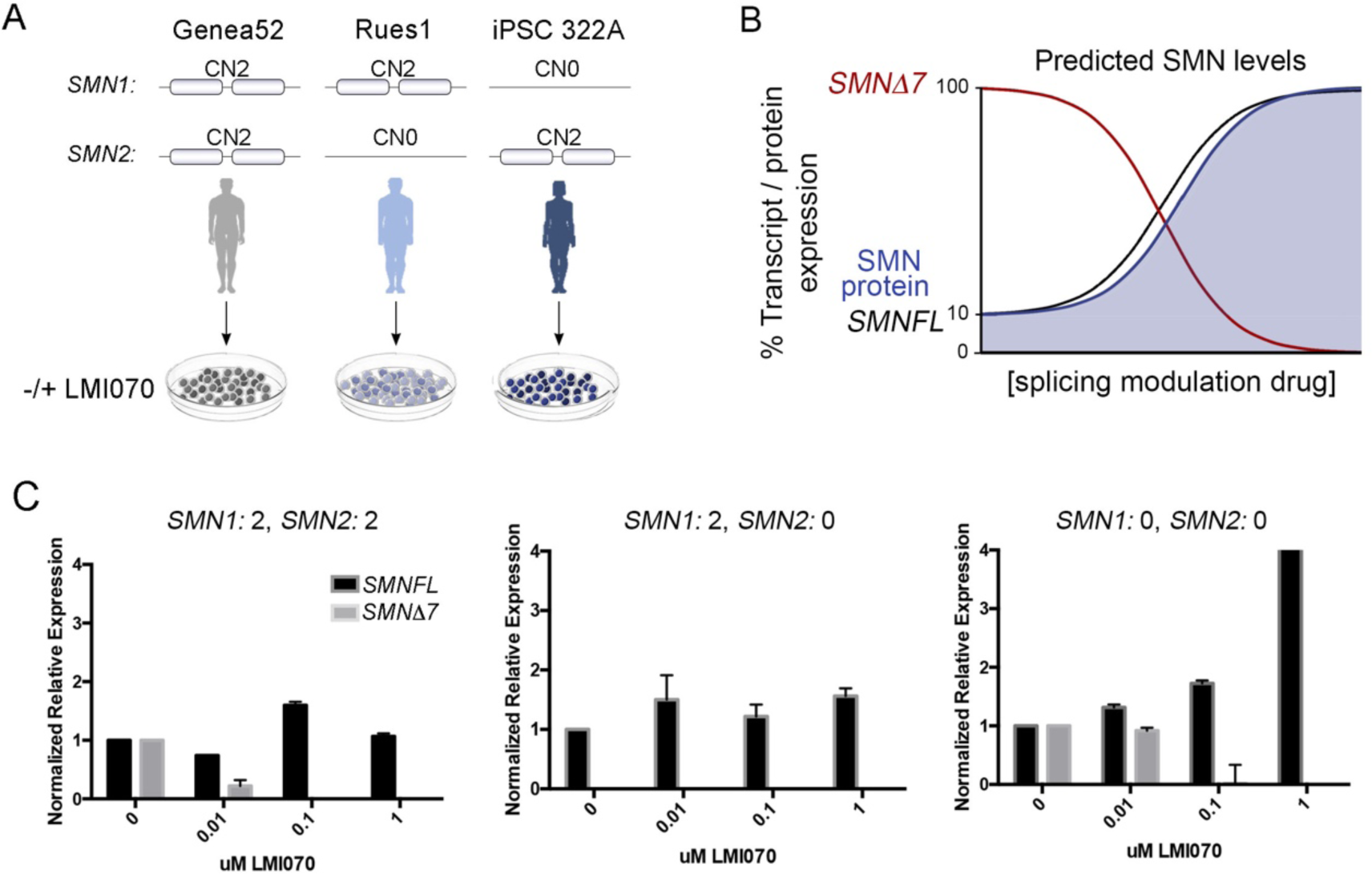
Characterization of LMI070 effects on *SMN* RNA transcript expression in PSCs *in vitro*. A. Cell lines from individual donors of varying *SMN1* and *SMN2* gene copy numbers were chosen to find effective doses of LMI070 treatment in culture, for subsequent cell-village experiments. B. In this model, as levels of *SMNdelta7* transcripts are decreased by the activity of LMI070, we expect a corresponding increase in levels of *SMN* full-length transcript and SMN protein. C. Measurements (by RT-PCR) of the levels of *SMN* full length (black) and *SMNdelta7* (grey) RNA transcripts in donor cell lines of varying *SMN* copy number. Note that *SMNdelta7* transcripts are produced only in the two donors with *SMN2* genes (left and right panels), and that the expression of such transcripts is reduced by treatment with LMI070.

**Figure S5.**
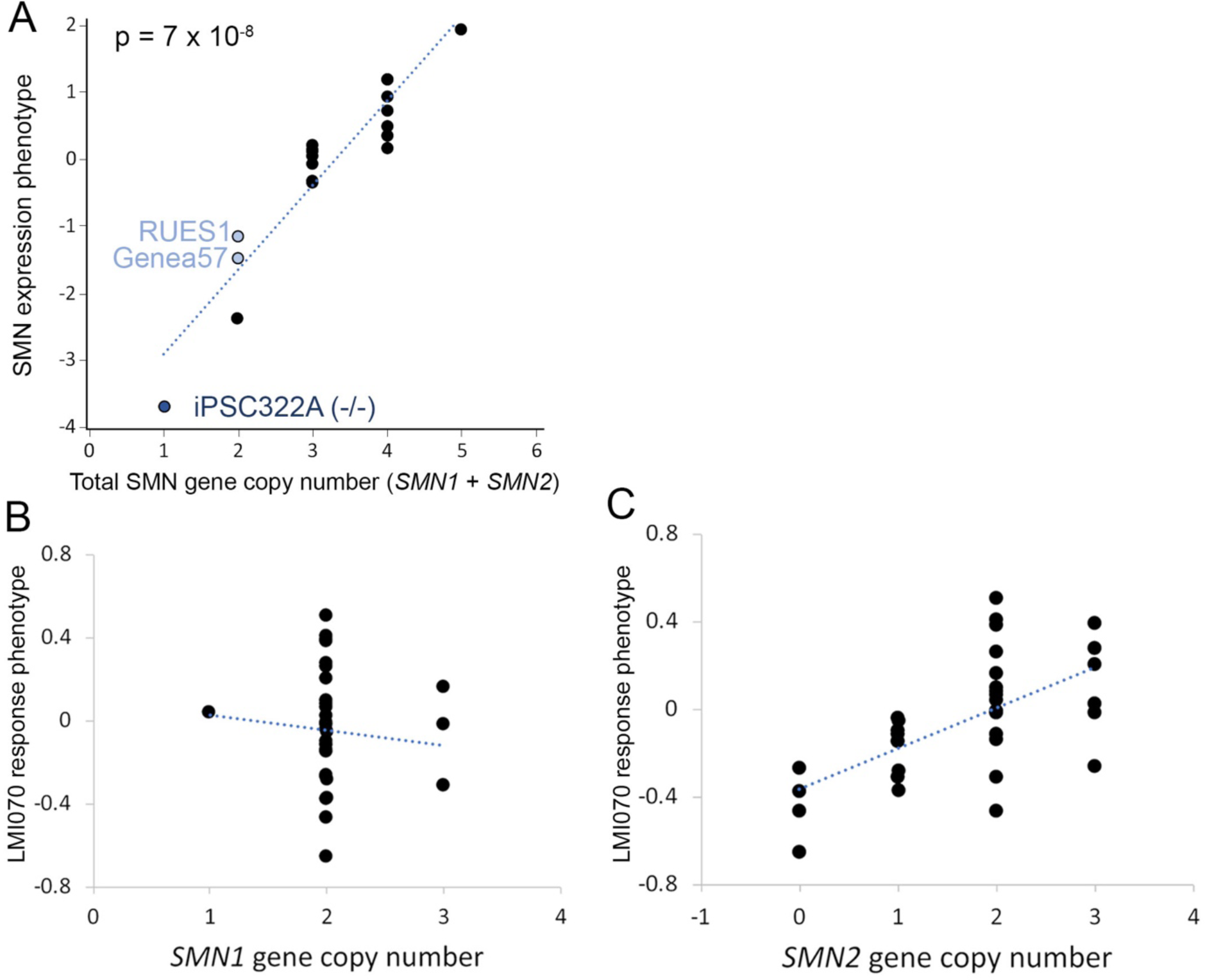
Using Census-seq to analyze genetic contributions to SMN protein phenotypes in a pilot village of hESCs. Companion replication data to Figures 5C and 6EF, drawing upon an additional cell village of hESC lines. A. Correlation of the Census-seq SMN protein-expression phenotype with common variation in the total (summed) copy number of *SMN1* and *SMN2* genes. B. Correlation of the Census-seq LMI070-response phenotype with *SMN1* gene copy number. C. Correlation of the Census-seq LMI070-response phenotype with *SMN2* gene copy number.

**Figure S6.**
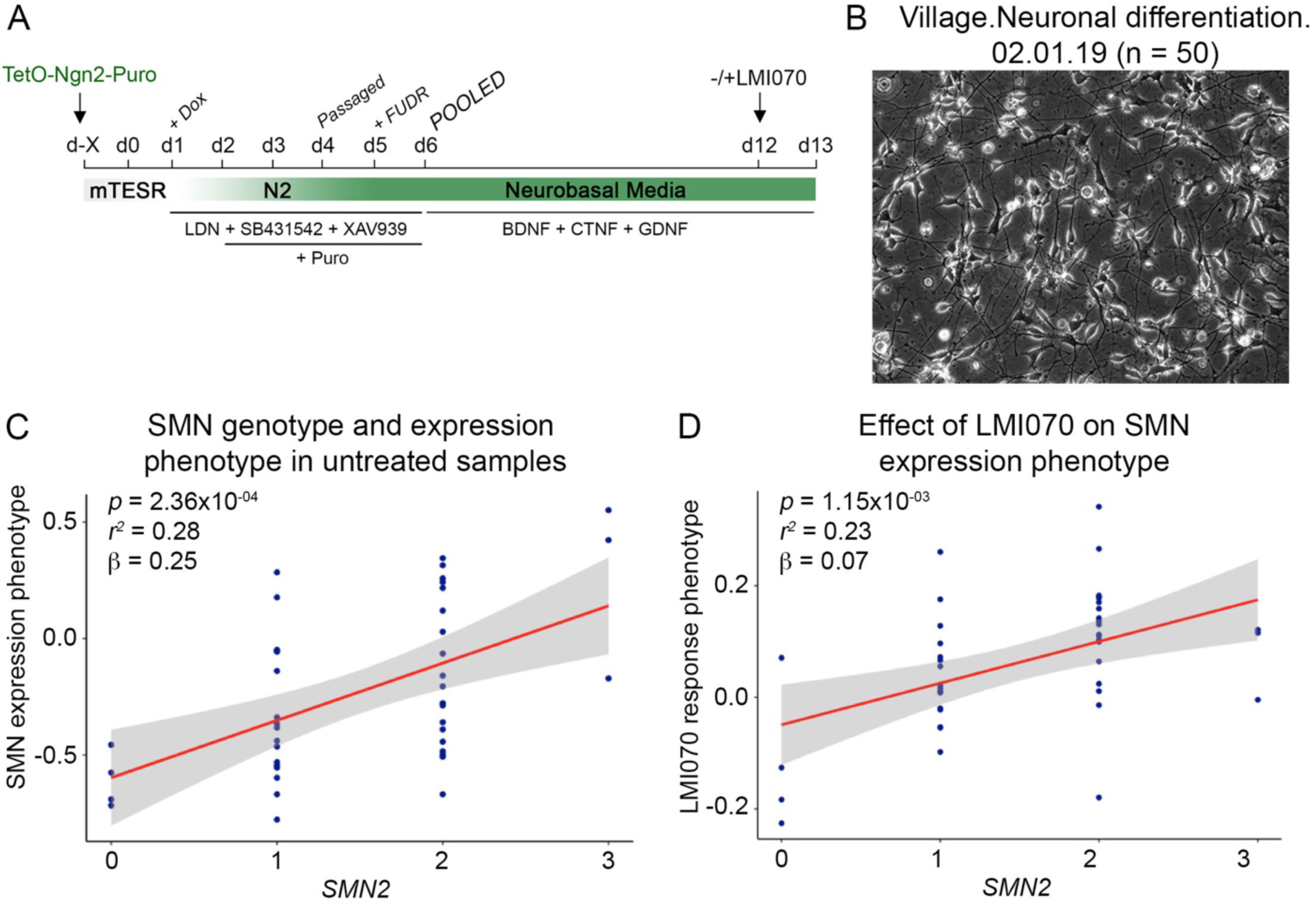
Pharmacogenetic analysis of response to LMI070 treatment in a village of differentiated neurons. A. Differentiation protocol used to generate a village of neurons derived *in vitro* from 50 iPSC donors. Cell lines transduced with lentivirus to drive *Ngn2* expression were induced with doxycycline, selected using puromycin, and pooled at day 6 of differentiation. The village was treated with LMI070 on day 12 and harvested for flow cytometry 24 hours later. B. Micrograph of the neuronal village at day 13 of differentiation. C. Correlation of the Census-seq SMN protein-expression phenotype with common variation in *SMN2* gene copy number. D. Correlation of the Census-seq LMI070 drug-response phenotype with common variation in *SMN2* gene copy number.

**Figure S7.**
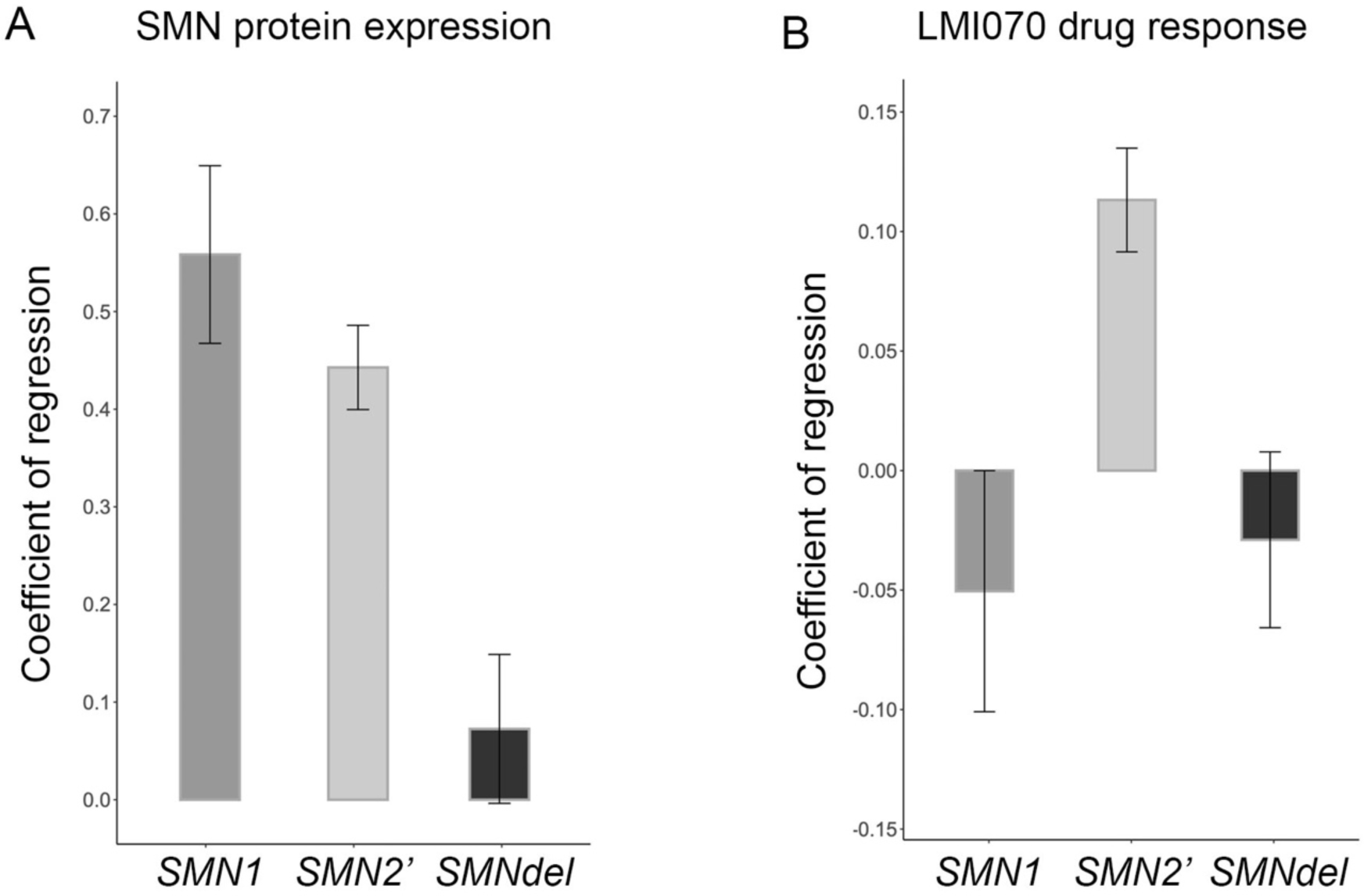
Contributions of *SMN1*, intact *SMN2* genes (excluding *SMNdel*) and *SMNdel* alleles to Census-seq phenotypes for SMN protein expression and response to splicing correction by LMI070. A. Contributions of *SMN1* genes, intact *SMN2* genes (*SMN2’*, which excludes *SMNdel* alleles), and *SMNdel* alleles to the SMN protein-expression Census-seq phenotype. Bars indicate coefficients of a linear regression of the SMN protein-expression phenotype against *SMN1, SMN2’*, and *SMNdel* gene copy numbers. Error bars indicate standard error. B. Contributions of *SMN1* genes, intact *SMN2* genes (*SMN2’*, which excludes *SMNdel* alleles), and *SMNdel* alleles to the LMI070-response Census-seq phenotype. Bars indicate coefficients of a linear regression of the SMN protein-expression phenotype against *SMN1, SMN2’*, and *SMNdel* gene copy numbers. Error bars indicate standard error.

## STAR Methods

### RESOURCE AVAILABILITY

#### Lead contact and materials availability

Further information and requests for resources and reagents should be directed to the Lead Contact, Steve McCarroll.

#### Data and code availability

All data reported in this manuscript, including Census-sequencing data, drug sensitivity measures, and other cell line features used in the analysis, will be available on a companion web site for the paper.

Custom code used in the analysis, and for generating all figures, is available at: https://github.com/broadinstitute/Drop-seq

SMN genotype-phenotype analyses: https://github.com/broadinstitute/dropseqrna/blob/master/transcriptome/R/ghoshs/Census_GenoPheno_Regression.R

### EXPERIMENTAL MODEL AND SUBJECT DETAILS

#### Acquisition of human pluripotent cell lines

As a source for human embryonic stem cells used in this study, we utilized a collection of curated hESCs we described previously (Merkle et al., 2017). Briefly, we sought to acquire a large cohort of hESCs approved for use by the National Institutes of Health (NIH) that were readily available and free of karyotypic or disease-causing abnormalities (http://grants.nih.gov/stem_cells/registry/current.htm). These cell lines were subjected to whole exome sequencing and whole genome sequencing, and were banked after minimal passaging (**Table S1, S3**).

To analyze cells from a larger number of individuals, using lines that could be readily acquired by any lab, we extended our analysis to include iPSCs from the cellular resource developed by the California Institute for Regenerative Medicine (CIRM). Cell lines of low passage number were obtained, expanded, submitted for whole genome sequencing, banked after minimal passaging, and regularly monitored for karyotypic abnormalities using the Illumina Global Screening Array (GSA). As a general rule, cell lines used in this study were passaged a maximum of 6 times after thawing to minimize the acquisition of karyotypic abnormalities or other deleterious mutations (**Table S1, S3)**.

## METHOD DETAILS

### Cell culture of Human pluripotent stem cells

Human embryonic stem cells and human induced pluripotent stem cells (hESCs and iPSCs, **Table S1**) were grown using StemFlex™ Medium (A3349401, Gibco™) supplemented with Normocin™ Antimicrobial Reagent (Invivogen, Ant-nr-1) on tissue culture dishes coated with Geltrex™ LDEV-Free Reduced Growth Factor Basement Membrane Matrix (A1413202, Gibco™) and maintained in 5% CO_2_ incubators at 37 °C. iPSCs were dissociated to single cell suspension using Accutase® (AT 104, Innovative Cell Technologies, Inc.). To prevent cell death, cell culture media was supplemented with ROCK inhibitor (Sigma, Y-27632, 10 µM) for 24hr following passaging. Cell lines were expanded to 10 cm plates, and when approaching confluency, cell lines were cryopreserved in CryoStor® (07930, StemCell Technologies) for further use. Cell lines were passaged a maximum of 6 times in an effort to preserve karyotypic integrity.

### Lentiviral transduction and neuronal differentiation

#### Lentiviral transduction

Cell lines were transduced with lentivirus cocktail to express tetracycline-inducible murine Neurogenin 2 (Ngn2) tagged with a puromycin resistance gene, in combination with tetracycline inducible GFP, as described (Ho et al., 2016; Nehme et al., 2018). PSCs were grown using StemFlex™ Medium (Gibco™, A3349401) on tissue culture dishes coated with Geltrex™ LDEV-Free Reduced Growth Factor Basement Membrane Matrix (Gibco™, A1413202) and maintained in 5% CO_2_ incubators at 37 °C. iPSCs were dissociated to single cell suspension using Accutase (Innovative Cell Technologies, Inc., AT 104). The concentration of cells in suspension was estimated using the Scepter Automated Cell Counter (Millipore Sigma). Cells were plated at a density of 100 000 cells/cm^2^ and incubated while in suspension with media containing 10 uM ROCK-Inhibitor (Sigma, Y27632), and lentivirus particles at a final MOI of 2. Three independent viruses were co-transduced to induce the expression of NGN2 using doxycycline (pTet-O-Ngn2-puro; pTet-O-EGFP; Ub-rtTA, gift from Marius Wernig; Lentivirus was produced by ALSTEM). Transduced cells were expanded to 10 cm plates and 10 vials (2M cells/vial) of each cell line was cryopreserved in CryoStor® (StemCell Technologies, 07930) for further use.

#### Neuronal differentiation

Differentiation of PSCs towards neuronal identity was performed as described (Nehme et al., 2018). Neuronal differentiation was initiated (day 1) by the addition of Doxycycline hyclate (2 µg/mL) to N2 supplemented media (Thermo Fisher, 17502048,) in the presence of patterning factors SB431542 (Tocris, 1614, 10 µM), XAV939 (Stemgent, 04-00046, 2 µM) and LDN-193189 (Stemgent, 04-0074, 100 nM) (referred to here as differentiation media). Non-transduced cells were removed from the cultures using Puromycin selection (5 µg/µL), from days 2 to 6. On day 3, cells were passaged into differentiation media supplemented with 5-Ethynyl-2’-deoxyuridine (Life Technologies, A10044, 10 µM). On day 6 of differentiation, cells were dissociated with Accutase®, counted using the Scepter™ Automated Cell Counter (Millipore Sigma), and villages of immature neurons were generated using 0.5 x 10^6^ cells from each donor cell line. Villages were plated at a density of 40 000 cells/cm^2^ in Neurobasal media (Gibco, 21103049) supplemented with B27 (Gibco, 17504044, 50X), doxycycline (2 µg/mL), brain-derived neurotrophic factor (BDNF), ciliary neurotrophic factor (CTNF), and glial cell-derived neurotrophic factor (GDNF) (R&D Systems 248-BD/CF, 257-NT/CF, and 212-GD/CF at 10 ng/mL each). Neuronal villages were co-cultured with murine glial cells (at a density of 70 000 cells/cm^2^) derived from postnatal pups (P0) as previously described (Di Giorgio et al., 2008). Villages were harvested at weekly intervals for Census-seq analysis. Neuronal villages were treated with LMI070 (see below) for 24 hours at day 12 of differentiation (**Figure S6**).

### Treatment with LMI070

To enhance splicing of the *SMN2* transcript to include exon 7, the small molecule LMI070 (also referred to as NVS-SM1 or Branaplam (Novartis; MedChemExpress, HY-19620)) was added to cell culture media for 24 hr under standard culture conditions. To determine the optimal dose of LMI070 in our cellular systems, a dose response was performed using a range of concentrations from 0.01 to 1 uM (**Figure S4**). Inclusion of exon 7 in the *SMN2* transcript was used as an indicator of effectivity. 0.1 uM was found to be optimal and used in all future experiments.

### Generation of villages of human pluripotent cell lines

Individual human pluripotent cell lines (cultured as described above) were dissociated using Accutase and resuspended in StemFlex supplemented with ROCK inhibitor. Approximately 0.5 x 10^6^ cells/cell line were combined to generate villages ranging from 10 to 115 donors and cells were plated at a density of 30 000 cells/cm^2^. To monitor donor composition over time in culture, 0.5 × 10^6^ cells were harvested during routine weekly passaging and samples were pelleted and stored at -20°C. Three different methodologies were used to quantify cellular concentrations depending on downstream applications (**Figure S1BCDE**).

### Quantification of cellular suspensions using the Countess™ hemocytometer

Cell enumeration using the Countess™ II Automated Cell Counter (AMQAX1000, Thermo Fisher) was used for routine cell density and viability estimations. Cell pellets generated from cell lines cultured in 6-well plate format were resuspended in 0.5 ml StemFlex supplemented with 10 uM ROCK inhibitor. 10 ul of a 1:1 mixture of cellular suspension and Trypan Blue solution (15250061, Thermo Fisher) was loaded into a Countess cell counting chambered slide and cell concentrations and viabilities were measured. To generate villages, 0.5 × 10^6^ viable cells from each donor cell line were mixed, 0.5 × 10^6^ cells of the village were sampled for downstream Census-seq analysis, and remaining cells were plated at a density of 30 000 cells/cm^2^.

### Quantification of cellular suspensions using the Scepter™ Automated Cell Counter

To generate villages of cells (including iPSCs and neural precursors), concentrations of cell suspensions were determined using the Scepter™ 2.0 Handheld Automated Cell Counter (Millipore Sigma, PHCC20060) equipped with 60 uM Scepter™ Cell Counting Sensors (Millipore Sigma, PHCC60050). Dilutions of cell suspensions (100 fold) were routinely quantified and concentrations indirectly calculated.

### Antibody-mediated pluripotency selection and precise sorting of cell lines using flow cytometry

To generate villages of pluripotent stem cells used to characterize growth/proliferation phenotypes (**Figure 2**), flow cytometry using antibodies selecting for markers of pluripotency was used to select for cells of the highest quality combined with precision counting. Prior to immunostaining, cells were dissociated using Accutase® and cell densities were measured using the Countess™ Hemocytometer. 5 × 10^6^ cells from each cell line were resuspended in FACS buffer (1 x PBS, 7.5% BSA, 1 mM EDTA, 10 uM Y27632, and Nomrocin™) and cells were immunostained with fluorescently labeled antibodies directed against SSEA4 and TRA-1-60 (BD Biosciences, Alexa Fluor® 647 Mouse anti-SSEA-4 Clone MC813-70 (RUO), 560219; Alexa Fluor® 488 Mouse anti-Human TRA-1-60 Antigen Clone TRA-1-60 (RUO), 560173) at a concentration of 1:500 for 30 min at RT. Cells were washed with FACS buffer repeatedly and stained with DAPI (1 ug/ml). 0.5 × 10^6^ DAPI-/SSEA4+/TRA-1-60+ cells were collected for each cell line using the BD FACSAria™ II flow cytometer (BD Biosciences). Collected cells were pooled and plated at a density of 30 000 /cm^2^.

### Phenotype-based sorting of SMN-high and SMN-low cellular populations

To allow for genotype-phenotype correlations based on SMN protein expression, villages of iPSCs or NGN2-induced neurons (at 14 days of differentiation) were immunostained with a monoclonal antibody directed against SMN (BD Biosciences, 610647) and segregated into new ‘sub-villages’ comprising populations of the lowest and highest SMN expressing cells.

Following dissociation with Accutase®, 10 × 10^6^ cells from villages of iPSCs or neurons were fixed and permeabilized using the Fixation/Permeabilization Solution Kit (BD Biosciences, 554714). Cells were resuspended in FACS buffer (1 x PBS, 7.5% BSA, 1 mM EDTA, 10 uM Y27632, and Nomrocin™) and incubated in the presence of anti-SMN at a concentration of 1: 500 for 30 min at RT. Cells were washed repeatedly in FACS buffer and incubated with donkey-anti-mouse-Alexa-647 secondary antibody (Life Technologies, A-31571) at a concentration of 1:5000 in FACS buffer supplemented with DAPI (1 ug/ml) for 30 min at RT. After repeated washing, cells were sorted using the BD FACSAria™ II flow cytometer (BD Biosciences).

Gating strategies were developed to select for single, live, intact cells in the absence of autofluorescence (using an empty 555 nm channel as control) (Figure S5). For neurons, an additional GFP (488 nm) selection gate was applied to separate Human neurons (GFP+) from co-cultured mouse glia. Cells determined to be positively stained for SMN (647 nm) were compared to unstained or secondary antibody-only controls, and gates were set using the patient cell line iPSC3222A as an indicator of low levels of SMN expression. To generate sub-villages, fractions encompassing the bottom (SMN-low) and top (SMN-high) 20% of cells were independently collected (500 000 cells each). An aliquot of unsorted cells representing donor representation of the original village was collected to serve as a control. Collected cells were pelleted and stored at -20°C for DNA isolation.

### Quantitative analysis of *SMN1* and *SMN2* transcript abundance by qPCR

SMN transcript abundance was quantified in cell lines of varying *SMN1* and *SMN2* copy number as follows: Genea52 (copy number *SMN1*:*SMN2*; 2/2), RUES1 (2/0), and iPSC322A (0/2). Cells were treated in the presence or absence of LMI070 as described. Total RNA was isolated from cell pellets using the RNeasy Mini Kit (Qiagen, 74104). cDNA was synthesized using iScript cDNA Synthesis Kit (Bio-Rad, 1708890). Two primer sets were used to detect *SMN* transcripts as described previously (Integrated DNA Technologies (Sumner et al., 2006): *SMNFL* spanning exons 6, 7 and 8 (forward, 5’-CAAAAAGAAGGAAGGTGCTCACATT-3’; reverse, 5’-GTGTCATTTAGTGCTGCTCTATGC-3’; probe, 5’-6FAM-CAGCATTTCTCCTTAATTTA-MGBNFQ-3’), and *SMNdelta7* spanning the exon 6-8 junction (forward, 5’-CATGGTACATGAGTGGCTATCATACTG-3’; reverse, 5’-TGGTGTCATTTAGTGCTGCTCTATG-3’; probe, 5’-6FAM-CCAGCATTTCCATATAATAGC-MGBNFQ-3’). mRNA levels were normalized to an internal control (Human Beta Glucuronidase (GUSB), Life Technologies, 4333767T). For samples treated with LMI070, *SMN1* and *SMN2* levels were compared to a DMSO-treated control. qRT-PCR was performed using TaqMan™ Fast Advanced Master Mix (ThermoFisher Scientific, 4444963).

### DNA Isolation from fixed or frozen cell pellets

Cell pellets were resuspended in 600 ul Cell Lysis Solution (Qiagen, 158906) supplemented with 3 ul Proteinase K (NEB, P8107S). Samples were incubated for 1 hr at 56°C. For samples that had been fixed with aldehydes (for example, for immunostaining prior to FACS), samples were further incubated overnight at 65°C. RNases were removed by incubating samples in the presence of 2 ul of RNase A (Qiagen, 158922) for 30 min at 37°C and samples were chilled for 5 on ice. Proteases were precipitated using 200 ul of Protein Precipitation Solution (Qiagen, 158910) and centrifuged for 10 min at 12 000 *g* at 4°C. DNA was precipitated from supernatants by adding 600 ul 100% isopropanol and centrifuging for 10 min at 12 000 *g* at 4°C. DNA pellets were washed with 600 ul 70% ethanol, and centrifuged for 5 min at 12 000 *g* at 4°C. Pellets were air dried and resuspended in 50 ul of dH20 or 10 mM Tris-Cl, pH 8.5.

### Generation and sequencing of libraries

Sequencing libraries were generated from isolated DNA using either TruSeq Nano DNA Library Prep Kit (Illumina, NP-101-1001) or Nextera DNA Flex Library Prep Kit (Illumina FC-121-1030) in combination with the NeoPrep Library Prep System (Illumina, SE-601-1001). Libraries were sequenced using the NextSeq 500 Sequencing System (Illumina, SY-415-1001) with the NextSeq 500 High Output v2 Kit (75 cycles, FC-404-2005). Runs were set up as a single 85 bp reads and included an index read when libraries were pooled. For scaled analyses, 16 Census-Seq samples are pooled into one NextSeq run with a minimum requirement of 16-32 million reads per library (about 1X coverage).

### Sequence Alignment Protocol

Raw sequence data were demultiplexed using the Picard tools ExtractIlluminaBarcodes and IlluminaBasecallsToSam. The resulting demultiplexed libraries were validated for both relative library size (library balance) as well as absolute size, to flag potential bioinformatic issues with demultiplexing, as well as benchside library generation issues. The demultiplexed libraries were then aligned to a human reference genome with BWA. The reference genome used was selected to match the same build used in the VCF file that contained the reference genotypes for the experiment’s donor pool.

For experiments for which human cells were grown on a bed of mouse glia, alignment was performed using a multi-organism reference and the reads were competitively aligned to both genomes. Sequencing reads were then filtered to reads that mapped at high quality (MQ>=10) to the human genome.

### Variant Call Format (VCF) pre-analysis Processing

Prior to running Census-seq, VCF files were processed to filter variants and add additional site-level information. Variants were first normalized to their appropriate reference sequence using BCFTools; this splits multiallelic SNPs into multiple biallelic SNPs, and sets the reference allele to be the reference base of the genome at that position. Variants that were monomorphic were dropped, as well as those without a PASS filter, where the site was flagged as problematic during VCF generation. Sites without rsID annotations were updated using information from dbSNP when possible, and otherwise site names were changed to chromosome:position:ref_allele:alt_allele. Allele frequencies calculated from the 1000 Genomes Project were annotated at all available sites.

### Computational analysis methods

Methods to detect the presence of individuals’ DNA within DNA mixtures have been a lively area of computational investigation (Egeland et al., 2003, Hu and Fung, 2003, Balding, 2003, Clayton et al., 1998). Our goal with Census-seq, Roll Call and CSI was to develop a suite of algorithms with which to detect and precisely quantify individual donors’ contributions to cell/DNA mixtures and to detect the presence of contaminating DNA/cells of known or unknown genotypes.

### Census-seq algorithm (precise quantification of donor contribution to cell/DNA mixtures)

The goal of Census-seq is to measure the contribution of each donor to a cell mixture – both to monitor population dynamics, and for quantitative phenotyping. We do this systematically, routinely and inexpensively, without the need for single-cell analysis, simply by lightly sequencing genomic DNA from the cells. The donor mixture determines what ratio of alleles are present at every SNP. We developed a gradient-descent algorithm to find the donor-mixing coefficients that maximize the likelihood of any observed sequence data.

For Census-seq to perform accurately, input sequencing and VCF data is filtered on a per-run basis. Sequence reads are filtered to high quality mappings (MQ>=10) on the autosomes that have not been flagged as PCR duplicates. VCF sites are considered if they meet all of the following criteria: each site has GQ score of at least 30, is a diploid site, is polymorphic in the subset of donors in the population, and at least 90% of donors have a genotype quality score >=30. In addition, for genotype array-based data where site quality scores may not be available, sites where the reference base is ambiguous [A/T, C/G] are not considered. Variant sites are also rejected if they are not common in the population – only sites with ∼5% allele frequency are included in analysis.

Given these filtered inputs, a matrix of donor genotypes and the counts of the reference and alternate allele at each variant are generated. Census-seq then uses these to find a vector of donor-specific contributions (to the mixture) that best explains the observed counts of alleles at each site in the sequence data. The algorithm initializes with the donor proportions set to equal values (1/number of donors), then runs through an estimation maximization (EM) procedure. During each step, the allele frequency of each site is calculated from the genotypes of the donors and their relative proportion in the pool. The initial likelihood of the sequencing data given the starting donor ratios is calculated at each SNP by the likelihood function (see below) and the results summed across all sites. To determine how to change the donor ratios to explain the data, an adjustment term is calculated for every donor/site, and the results are summed across sites for each donor. This adjustment factor is then scaled by an additional parameter and added to each donor’s representation. To determine this scaling value the algorithm employs a univariate optimizer to maximize the donor likelihood. The adjustment is then applied to the data, and the algorithm repeats the adjustment/ likelihood optimization loop until convergence.

Log-likelihood Function:

For any set of donor-mixing coefficients (and resulting allele frequencies in the DNA mixture), he log-likelihood of the Census-seq sequencing data is calculated as:

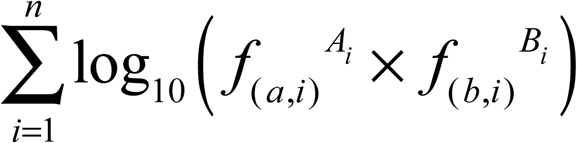

where *i* (= 1, 2, …, n) indexes the full set of SNPs used in analysis; *f*_*(a,i)*_ and *f*_*(b,i)*_ are the frequencies of the two alleles for SNP *i*; and *A*_*i*_ and *B*_*i*_ are the numbers of observations of these alleles in the Census-seq sequencing data.

The derivative of the above likelihood function with respect to the donor-specific mixing coefficients is used to calculate a gradient ascent direction in the form of an adjustment factor for each donor; this adjustment term reflects the extent to which an increase (or decrease) in that donor’s mixing coefficient improved the likelihood of the Census-seq data. During each loop of the EM, each donor’s mixing coefficient (contribution estimate) is adjusted by a factor that is proportional to the result of the following formula:

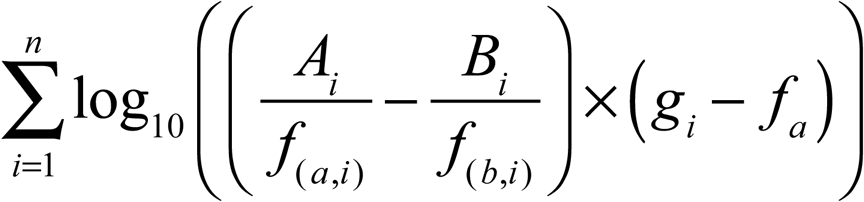

Where *i* (= 1, 2, …, n) indexes the full set of SNPs used in analysis; *g*_*i*_ is the genotype of the donor at SNP *i* (= 1 if homozygous for the reference allele, 0 if homozygous for the alternate allele, and 0.5 if heterozygous); and *f*_*(a,i)*_ and *f*_*(b,i)*_ are the frequencies of the two alleles for SNP *i*.

### Roll Call algorithm (to establish which donors have contributed to a cell/DNA mixture)

Census-seq requires a complete and accurate list of donors in order to estimate each donor’s contribution accurately. However, cell lines can become contaminated with cells from other donors; tubes can be mislabeled; and sample swaps can occur even despite best efforts. The probability of such errors increases with the number of cell lines in an experiment – thus, population-scale experiments are inherently more vulnerable to error. We thus developed an algorithm (“Roll Call”) to create a complete list of the donors who have contributed to a cell/DNA mixture (from a larger set of genomically characterized candidates, for example, all of the cell lines in a lab). Roll Call uses each donor’s private (singleton) alleles to find evidence that a donor is present in a mixture in which s/he doesn’t belong. To do this, we measure (for all of the sequence reads that touch these sites) the fraction of the sequence reads that individual’s IRVs. Since this donor is in principle the only source of alternate alleles at the IRV sites, the fraction of these alleles observed can be directly related to the presence of the donor to the mixture. In the equation below, we count the observations (sequence reads) that (in the absence of sequence errors) could only arise from a given individual, divided by the total number of reads at those sites. This is less precise than Census-seq at quantifying donor representation, because it draws upon a small fraction of all variable sites; the utility of Roll Call is to search through a very large set of potential donors to determine presence/absence and thereby authenticate a village prior to Census-seq analysis.

Roll Call uses the same VCF and sequencing read filtering as Census-seq, with one exception - instead of retaining common sites, Roll Call leverages IRVs (rare identifying variants) - sites that are private to a single donor in the VCF. Since these IRVs are the only source of alternate alleles in the sequencing data, the fraction of alleles in the sequencing data observed in a sequencing pool can directly be related to the proportion of donors in the pool.

To generate the counts of IRVs, the algorithm filters the VCF to a set of heterozygous sites that are private to donors in the VCF. The pileup of reference and alternate alleles is generated at those sites in the sequencing data. For each donor, those pileups are then aggregated into a single result, and the Roll Call score is calculated:

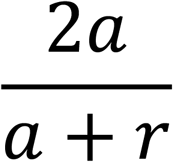

where *a* is the number of alternate alleles (and r is the number of reference alleles) observed for that donor’s IRVs at heterozygous sites.

Note that PCR and sequencing error cause the Roll Call score to be slightly positive even for donors who have not contributed to a mixture. For any experiment, a background null distribution of Roll Call scores can be estimated by including (in the input VCF) many individuals not expected to have contributed to the cell/DNA mixture; this distribution can then inform the selection of an experiment-specific threshold for the Roll Call score.

We routinely use Roll Call to validate and if necessary refine the donor list prior to Census-seq analysis.

### CSI - Contaminating Sample Identifier (detection of cells/DNA from genomically unknown donors)

What if a pool were visited by cells from a genomically “unknown” donor? If we don’t have genotypes for the contaminating donor(s) *a priori*, we need another way to detect the presence of “unknown unexpected” visitors. We developed the CSI algorithm to do this. CSI utilizes observations of alternate alleles that could not have arisen from the expected donors’ genomes and must therefore represent sequencing errors or contaminating cells. We first identify all genomic sites and alleles that are absent among the donors we expect in the pool but present at some minimum frequency in the wider population (as estimated from the Thousand Genomes Project or gnomAD data). We then look for evidence of such alleles in the village DNA sequencing data. By correcting for sequencing error rate, we distinguish between two models: sequencing errors and an unwelcome visitor.

CSI utilizes observations of alternate alleles that could not have arisen from the expected donors’ genomes and must therefore represent sequencing errors or contaminating cells. We first identify all alleles that are absent among the donors we expect in the pool but present at some minimum frequency in the wider population.

CSI uses the same VCF and sequencing read filtering as Census-Seq with one exception - the variant sites selected from the VCF are those for which all donors in the experiment have the reference genotype, and the minor allele frequency (MAF) of these sites in a wider population is at least 2.5% This population allele frequency can be computed from a variety of potential sources, including (i) all donors in the VCF file not expected to be present in the experimental mixture; or, (ii) an external reference population such as those provided by the 1000 Genomes Project or gnomAD.

To calculate the CSI intrusion score, we count observations of of reference and alternate alleles across all such sites and aggregate the results. We then calculate the CSI intrusion score by taking into account the sequencing error rate and average minor allele frequency of the sites queried.

CSI intrusion score:

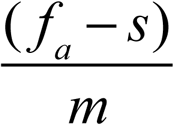

Where Fa is the fraction of sequencing reads (at these sites) observed to contain the alleles that are absent among the candidate donors; s is the sequencing error rate; and m is the mean minor allele frequency (of these alleles) in the population from which the potential donors are sampled. Note that while the CSI intrusion score is proportional to the degree of contamination from unexpected donor(s), two possibilities can cause it to deviate: (i) if the unexpected donor is related to an expected/candidate donor; or (ii) if the unexpected donor comes from a different population than the population used to estimate the mean MAF of the absent alleles. (However, *changes* in the CSI score across time are likely to represent changes in the contribution of a contaminating donor.) For this reason, we generally use the CSI score to authenticate or flag cell villages rather than to precisely measure contamination; a cell village contaminated by an unknown donor would not be suitable for Census-seq analysis anyway.

## QUANTIFICATION AND STATISTICAL ANALYSIS

In analyzing the SMN-expression phenotype, we aggregated results from two villages of iPSCs, one of 113 donors, which we called CIRM1, and another with 38 donors, which we called SMN6.

For each village, we calculate an SMN-expression phenotype measure for each donor as the log10-fold change in the representation of that donor in the SMN-high fraction of the village over their representation in the SMN-low fraction of the village: *SMNexpression = log*10 (*rep*_*high*_/*rep*_*low*_).

Census-seq has reduced precision in quantifying donors who have contributed less than 0.2% of the cells/DNA in a mixture. To account for this, as a quality control measure, we exclude from association analysis any donors who have less than 0.3% representation in any of the relevant derived villages (in this case, SMN-high or SMN-low). 72 of the donors from the village of 113 and 37 donors from the village of 38 donors passed this filtering step and were used in downstream genetic analysis.

20 of the 28 donors shared between the two villages were above threshold in both villages. To construct an aggregate table of the two villages, we averaged the SMN-expression phenotype measurements from both villages for these 20 donors. Then, for the remaining 76 unique donors, we used the SMN-expression phenotype calculated from the one village in which it passed QC.

This resulted in a set of 96 donors with SMN-expression phenotype measurements suitable for genetic analysis. We performed linear regression of these 96 measurements against the number of copies of the *SMN* genes (**Figure 4C**).

### Measurement of quantitative phenotype and of quantitative drug response

We had collected two additional fractions of SMN-high and SMN-low for each village after treatment with the drug LMI070. Using these two LMI070-treated fractions, we calculated *SMNexpression*for the donors in each village when treated with LMI070. We then calculated the difference of *SMNexpression* of the donor when treated with LMI070 and *SMNexpression*of the donor when not treated with LMI070, calling this difference *LMI*070*DrugResponse. LMI*070*DrugResponse* = *SMNexpression*_*LMI070treated*_ − *SMNexpression*_*control*_

Since we used two new fractions to calculate drug response, our quality control of the data had to adjust to take these into account. In particular, we filtered the data from each village to only include donors that had a representation of at least 0.3% in all 4 fractions. 67 donors from the village of 113 and 36 of the village of 38 passed QCl, with 11 of the 28 common donors passing in both villages. We averaged values for the 11 common donors and created an aggregate table to use for downstream analysis that had measurements for 92 unique donors. Finally, we ran a linear regression of the effect of *SMN2* copy number on LMI070 drug response and found that there was a significant linear correlation between *SMN2* copy number and LMI070 drug response (**Figure 6D**) that was even stronger when we discount any copies of the *SMNdel* variant of the gene in the regression model (**Figure 6E**).

## References

Adelmann, C. H., Wang, T., Sabatini, D. M. & Lander, E. S. 2019. Genome-Wide CRISPR/Cas9 Screening for Identification of Cancer Genes in Cell Lines. Methods Mol Biol, 1907, 125–136.

Balding, D. J. 2003. Likelihood-based inference for genetic correlation coefficients. Theor Popul Biol, 63, 221–30.

Boyle, E. A., Li, Y. I. & Pritchard, J. K. 2017. An Expanded View of Complex Traits: From Polygenic to Omnigenic. Cell, 169, 1177–1186.

Burnett, B. G., Munoz, E., Tandon, A., Kwon, D. Y., Sumner, C. J. & Fischbeck, K. H. 2009. Regulation of SMN protein stability. Mol Cell Biol, 29, 1107–15.

Cahan, P. & Daley, G. Q. 2013. Origins and implications of pluripotent stem cell variability and heterogeneity. Nat Rev Mol Cell Biol, 14, 357–68.

Canver, M. C., Smith, E. C., Sher, F., Pinello, L., Sanjana, N. E., Shalem, O., Chen, D. D., Schupp, P. G., Vinjamur, D. S., Garcia, S. P., Luc, S., Kurita, R., Nakamura, Y., Fujiwara, Y., Maeda, T., Yuan, G. C., Zhang, F., Orkin, S. H. & Bauer, D. E. 2015. BCL11A enhancer dissection by Cas9-mediated in situ saturating mutagenesis. Nature, 527, 192–7.

Chen, T. H. 2020. New and Developing Therapies in Spinal Muscular Atrophy: From Genotype to Phenotype to Treatment and Where Do We Stand? Int J Mol Sci, 21.

Cheung, A. K., Hurley, B., Kerrigan, R., Shu, L., Chin, D. N., Shen, Y., O’brien, G., Sung, M. J., Hou, Y., Axford, J., Cody, E., Sun, R., Fazal, A., Fridrich, C., Sanchez, C. C., Tomlinson, R. C., Jain, M., Deng, L., Hoffmaster, K., Song, C., Van Hoosear, M., Shin, Y., Servais, R., Towler, C., Hild, M., Curtis, D., Dietrich, W. F., Hamann, L. G., Briner, K., Chen, K. S., Kobayashi, D., Sivasankaran, R. & Dales, N. A. 2018. Discovery of Small Molecule Splicing Modulators of Survival Motor Neuron-2 (SMN2) for the Treatment of Spinal Muscular Atrophy (SMA). J Med Chem, 61, 11021–11036.

Cheung, V. G., Spielman, R. S., Ewens, K. G., Weber, T. M., Morley, M. & Burdick, J. T. 2005. Mapping determinants of human gene expression by regional and genomewide association. Nature, 437, 1365–9.

Clayton, T. M., Whitaker, J. P., Sparkes, R. & Gill, P. 1998. Analysis and interpretation of mixed forensic stains using DNA STR profiling. Forensic Sci Int, 91, 55–70.

Di Giorgio, F. P., Boulting, G. L., Bobrowicz, S. & Eggan, K. C. 2008. Human embryonic stem cell-derived motor neurons are sensitive to the toxic effect of glial cells carrying an ALS-causing mutation. Cell Stem Cell, 3, 637–48.

Egeland, T., Dalen, I. & Mostad, P. F. 2003. Estimating the number of contributors to a DNA profile. Int J Legal Med, 117, 271–5.

Finkel, R. S., Chiriboga, C. A., Vajsar, J., Day, J. W., Montes, J., De Vivo, D. C., Yamashita, M., Rigo, F., Hung, G., Schneider, E., Norris, D. A., Xia, S., Bennett, C. F. & Bishop, K. M. 2016. Treatment of infantile-onset spinal muscular atrophy with nusinersen: a phase 2, open-label, dose-escalation study. Lancet, 388, 3017–3026.

Gasperini, M., Hill, A. J., Mcfaline-Figueroa, J. L., Martin, B., Kim, S., Zhang, M. D., Jackson, D., Leith, A., Schreiber, J., Noble, W. S., Trapnell, C., Ahituv, N. & Shendure, J. 2019. A Genome-wide Framework for Mapping Gene Regulation via Cellular Genetic Screens. Cell, 176, 377–390 e19.

Groen, E. J. N., Talbot, K. & Gillingwater, T. H. 2018. Advances in therapy for spinal muscular atrophy: promises and challenges. Nat Rev Neurol, 14, 214–224.

Handsaker, R. E., Korn, J. M., Nemesh, J. & Mccarroll, S. A. 2011. Discovery and genotyping of genome structural polymorphism by sequencing on a population scale. Nat Genet, 43, 269–76.

Handsaker, R. E., Van Doren, V., Berman, J. R., Genovese, G., Kashin, S., Boettger, L. M. & Mccarroll, S. A. 2015. Large multiallelic copy number variations in humans. Nat Genet, 47, 296–303.

Hsu, J. Y., Fulco, C. P., Cole, M. A., Canver, M. C., Pellin, D., Sher, F., Farouni, R., Clement, K., Guo, J. A., Biasco, L., Orkin, S. H., Engreitz, J. M., Lander, E. S., Joung, J. K., Bauer, D. E. & Pinello, L. 2018. CRISPR-SURF: discovering regulatory elements by deconvolution of CRISPR tiling screen data. Nat Methods, 15, 992–993.

Hsu, P. D., Lander, E. S. & Zhang, F. 2014. Development and applications of CRISPR-Cas9 for genome engineering. Cell, 157, 1262–78.

Hu, Y. Q. & Fung, W. K. 2003. Interpreting DNA mixtures with the presence of relatives. Int J Legal Med, 117, 39–45.

Hua, Y., Sahashi, K., Hung, G., Rigo, F., Passini, M. A., Bennett, C. F. & Krainer, A. R. 2010. Antisense correction of SMN2 splicing in the CNS rescues necrosis in a type III SMA mouse model. Genes Dev, 24, 1634–44.

Hua, Y., Vickers, T. A., Okunola, H. L., Bennett, C. F. & Krainer, A. R. 2008. Antisense masking of an hnRNP A1/A2 intronic splicing silencer corrects SMN2 splicing in transgenic mice. Am J Hum Genet, 82, 834–48.

Khera, A. V., Chaffin, M., Aragam, K. G., Haas, M. E., Roselli, C., Choi, S. H., Natarajan, P., Lander, E. S., Lubitz, S. A., Ellinor, P. T. & Kathiresan, S. 2018. Genome-wide polygenic scores for common diseases identify individuals with risk equivalent to monogenic mutations. Nat Genet, 50, 1219–1224.

Kilpinen, H., Goncalves, A., Leha, A., Afzal, V., Alasoo, K., Ashford, S., Bala, S., Bensaddek, D., Casale, F. P., Culley, O. J., Danecek, P., Faulconbridge, A., Harrison, P. W., Kathuria, A., Mccarthy, D., Mccarthy, S. A., Meleckyte, R., Memari, Y., Moens, N., Soares, F., Mann, A., Streeter, I., Agu, C. A., Alderton, A., Nelson, R., Harper, S., Patel, M., White, A., Patel, S. R., Clarke, L., Halai, R., Kirton, C. M., Kolb-Kokocinski, A., Beales, P., Birney, E., Danovi, D., Lamond, A. I., Ouwehand, W. H., Vallier, L., Watt, F. M., Durbin, R., Stegle, O. & Gaffney, D. J. 2017. Common genetic variation drives molecular heterogeneity in human iPSCs. Nature, 546, 370–375.

Lam, M., Chen, C. Y., Li, Z., Martin, A. R., Bryois, J., Ma, X., Gaspar, H., Ikeda, M., Benyamin, B., Brown, B. C., Liu, R., Zhou, W., Guan, L., Kamatani, Y., Kim, S. W., Kubo, M., Kusumawardhani, A., Liu, C. M., Ma, H., Periyasamy, S., Takahashi, A., Xu, Z., Yu, H., Zhu, F., SCHIZOPHRENIA WORKING GROUP OF THE PSYCHIATRIC GENOMICS, C., Indonesia Schizophrenia, C., Genetic, R. O. S. N.-C., The, N., Chen, W. J., Faraone, S., Glatt, S. J., He, L., Hyman, S. E., Hwu, H. G., Mccarroll, S. A., Neale, B. M., Sklar, P., Wildenauer, D. B., Yu, X., Zhang, D., Mowry, B. J., Lee, J., Holmans, P., Xu, S., Sullivan, P. F., Ripke, S., O’Donovan, M. C., Daly, M. J., Qin, S., Sham, P., Iwata, N., Hong, K. S., Schwab, S. G., Yue, W., Tsuang, M., Liu, J., Ma, X., Kahn, R. S., Shi, Y. & Huang, H. 2019. Comparative genetic architectures of schizophrenia in East Asian and European populations. Nat Genet, 51, 1670–1678.

Larson, M. H., Gilbert, L. A., Wang, X., Lim, W. A., Weissman, J. S. & Qi, L. S. 2013. CRISPR interference (CRISPRi) for sequence-specific control of gene expression. Nat Protoc, 8, 2180–96.

Lefebvre, S., Burglen, L., Reboullet, S., Clermont, O., Burlet, P., Viollet, L., Benichou, B., Cruaud, C., Millasseau, P., Zeviani, M. & et al. 1995. Identification and characterization of a spinal muscular atrophy-determining gene. Cell, 80, 155–65.

Liang, G. & Zhang, Y. 2013. Genetic and epigenetic variations in iPSCs: potential causes and implications for application. Cell Stem Cell, 13, 149–59.

Lin, S. S., Delaura, S. & Jones, E. M. 2020. The CIRM iPSC repository. Stem Cell Res, 44, 101671.

Lo Sardo, V., Ferguson, W., Erikson, G. A., Topol, E. J., Baldwin, K. K. & Torkamani, A. 2017. Influence of donor age on induced pluripotent stem cells. Nat Biotechnol, 35, 69–74.

Loh, P. R., Genovese, G., Handsaker, R. E., Finucane, H. K., Reshef, Y. A., Palamara, P. F., Birmann, B. M., Talkowski, M. E., Bakhoum, S. F., Mccarroll, S. A. & Price, A. L. 2018. Insights into clonal haematopoiesis from 8,342 mosaic chromosomal alterations. Nature, 559, 350–355.

Lorson, C. L., Hahnen, E., Androphy, E. J. & Wirth, B. 1999. A single nucleotide in the SMN gene regulates splicing and is responsible for spinal muscular atrophy. Proc Natl Acad Sci U S A, 96, 6307–11.

Mcfarland, J. M., Paolella, B. R., Warren, A., Geiger-Schuller, K., Shibue, T., Rothberg, M., Kuksenko, O., Jones, A., Chambers, E., Dionne, D., Bender, S., Wolpin, B. W., Ghandi, M., Tirosh, I., Rozenblatt-Rosen, O., Roth, J. A., Tgolub, T. R., Regev, A., Aguirre, A. J., Vazquez, F. & Tsherniak, A. 2019. Multiplexed single-cell profiling of post-perturbation transcriptional responses to define cancer vulnerabilities and therapeutic mechanism of action. bioRxiv.

Merkle, F. T., Ghosh, S., Genovese, G., Handsaker, R. E., Seva Kashin, K., Karczewski Meyer, D., O’Dushlaine, C., Palotie, A., Pato, C., Pato, M., Schier, A. F., Macarthur, D., Mccarroll, S. A. & Eggan, K. in revision. Biological insights from the whole genome sequences of 143 widely available human embryonic stem cell lines. (In Revision).

Merkle, F. T., Ghosh, S., Kamitaki, N., Mitchell, J., Avior, Y., Mello, C., Kashin, S., Mekhoubad, S., Ilic, D., Charlton, M., Saphier, G., Handsaker, R. E., Genovese, G., Bar, S., Benvenisty, N., Mccarroll, S. A. & Eggan, K. 2017. Human pluripotent stem cells recurrently acquire and expand dominant negative P53 mutations. Nature, 545, 229–233.

Meyer, K., Marquis, J., Trub, J., Nlend Nlend, R., Verp, S., Ruepp, M. D., Imboden, H., Barde, I., Trono, D. & Schumperli, D. 2009. Rescue of a severe mouse model for spinal muscular atrophy by U7 snRNA-mediated splicing modulation. Hum Mol Genet, 18, 546–55.

Monani, U. R., Lorson, C. L., Parsons, D. W., Prior, T. W., Androphy, E. J., Burghes, A. H. & Mcpherson, J. D. 1999. A single nucleotide difference that alters splicing patterns distinguishes the SMA gene SMN1 from the copy gene SMN2. Hum Mol Genet, 8, 1177–83.

Morley, M., Molony, C. M., Weber, T. M., Devlin, J. L., Ewens, K. G., Spielman, R. & Cheung, V. G. 2004. Genetic analysis of genome-wide variation in human gene expression. Nature, 430, 743–7.

Nehme, R., Zuccaro, E., Ghosh, S. D., Li, C., Sherwood, J. L., Pietilainen, O., Barrett, L. E., Limone, F., Worringer, K. A., Kommineni, S., Zang, Y., Cacchiarelli, D., Meissner, A., Adolfsson, R., Haggarty, S., Madison, J., Muller, M., Arlotta, P., Fu, Z., Feng, G. & Eggan, K. 2018. Combining NGN2 Programming with Developmental Patterning Generates Human Excitatory Neurons with NMDAR-Mediated Synaptic Transmission. Cell Rep, 23, 2509–2523.

Neimark, J. 2015. Line of attack. Science, 347, 938–40.

Nelson-Rees, W. A., Daniels, D. W. & Flandermeyer, R. R. 1981. Cross-contamination of cells in culture. Science, 212, 446–52.

Nishizawa, M., Chonabayashi, K., Nomura, M., Tanaka, A., Nakamura, M., Inagaki, A., Nishikawa, M., Takei, I., Oishi, A., Tanabe, K., Ohnuki, M., Yokota, H., Koyanagi-Aoi, M., Okita, K., Watanabe, A., Takaori-Kondo, A., Yamanaka, S. & Yoshida, Y. 2016. Epigenetic Variation between Human Induced Pluripotent Stem Cell Lines Is an Indicator of Differentiation Capacity. Cell Stem Cell, 19, 341–54.

Palacino, J., Swalley, S. E., Song, C., Cheung, A. K., Shu, L., Zhang, X., Van Hoosear, M., Shin, Y., Chin, D. N., Keller, C. G., Beibel, M., Renaud, N. A., Smith, T. M., Salcius, M., Shi, X., Hild, M., Servais, R., Jain, M., Deng, L., Bullock, C., Mclellan, M., Schuierer, S., Murphy, L., Blommers, M. J., Blaustein, C., Berenshteyn, F., Lacoste, A., Thomas, J. R., Roma, G., Michaud, G. A., Tseng, B. S., Porter, J. A., Myer, V. E., Tallarico, J. A., Hamann, L. G., Curtis, D., Fishman, M. C., Dietrich, W. F., Dales, N. A. & Sivasankaran, R. 2015. SMN2 splice modulators enhance U1-pre-mRNA association and rescue SMA mice. Nat Chem Biol, 11, 511–7.

Pickrell, J. K., Marioni, J. C., Pai, A. A., Degner, J. F., Engelhardt, B. E., Nkadori, E., Veyrieras, J. B., Stephens, M., Gilad, Y. & Pritchard, J. K. 2010. Understanding mechanisms underlying human gene expression variation with RNA sequencing. Nature, 464, 768–72.

Ramdas, S. & Servais, L. 2020. New treatments in spinal muscular atrophy: an overview of currently available data. Expert Opin Pharmacother, 21, 307–315.

Rouhani, F., Kumasaka, N., De Brito, M. C., Bradley, A., Vallier, L. & Gaffney, D. 2014. Genetic background drives transcriptional variation in human induced pluripotent stem cells. PLoS Genet, 10, e1004432.

SCHIZOPHRENIA WORKING GROUP OF THE PSYCHIATRIC GENOMICS, C. 2014. Biological insights from 108 schizophrenia-associated genetic loci. Nature, 511, 421–7.

Shalem, O., Sanjana, N. E., Hartenian, E., Shi, X., Scott, D. A., Mikkelson, T., Heckl, D., Ebert, B. L., Root, D. E., Doench, J. G. & Zhang, F. 2014. Genomescale CRISPR-Cas9 knockout screening in human cells. Science, 343, 84–87.

Singh, R. N. & Singh, N. N. 2018. Mechanism of Splicing Regulation of Spinal Muscular Atrophy Genes. Adv Neurobiol, 20, 31–61.

Stranger, B. E., Forrest, M. S., Dunning, M., Ingle, C. E., Beazley, C., Thorne, N., Redon, R., Bird, C. P., De Grassi, A., Lee, C., Tyler-Smith, C., Carter, N., Scherer, S. W., Tavare, S., Deloukas, P., Hurles, M. E. & Dermitzakis, E. 2007a. Relative impact of nucleotide and copy number variation on gene expression phenotypes. Science, 315, 848–53.

Stranger, B. E., Nica, A. C., Forrest, M. S., Dimas, A., Bird, C. P., Beazley, C., Ingle, C. E., Dunning, M., Flicek, P., Koller, D., Montgomery, S., Tavare, S., Deloukas, P. & Dermitzakis, E. T. 2007b. Population genomics of human gene expression. Nat Genet, 39, 1217–24.

Sumner, C. J., Kolb, S. J., Harmison, G. G., Jeffries, N. O., Schadt, K., Finkel, R. S., Dreyfuss, G. & Fischbeck, K. H. 2006. SMN mRNA and protein levels in peripheral blood: biomarkers for SMA clinical trials. Neurology, 66, 1067–73.

Vijzelaar, R., Snetselaar, R., Clausen, M., Mason, A. G., Rinsma, M., Zegers, M., Molleman, N., Boschloo, R., Yilmaz, R., Kuilboer, R., Lens, S., Sulchan, S. & Schouten, J. 2019. The frequency of SMN gene variants lacking exon 7 and 8 is highly population dependent. PLoS One, 14, e0220211.

Wang, T., Wei, J. J., Sabatini, D. M. & Lander, E. S. 2014. Genetic screens in human cells using the CRISPR-Cas9 system. Science, 343, 80–4.

Yu, C., Mannan, A. M., Yvone, G. M., Ross, K. N., Zhang, Y. L., Marton, M. A., Taylor, B. R., Crenshaw, A., Gould, J. Z., Tamayo, P., Weir, B. A., Tsherniak, A., Wong, B., Garraway, L. A., Shamji, A. F., Palmer, M. A., Foley, M. A., Winckler, W., Schreiber, S. L., Kung, A. L. & Golub, T. R. 2016. High-throughput identification of genotype-specific cancer vulnerabilities in mixtures of barcoded tumor cell lines. Nat Biotechnol, 34, 419–23.

Zhang, Y., Pak, C., Han, Y., Ahlenius, H., Zhang, Z., Chanda, S., Marro, S., Patzke, C., Acuna, C., Covy, J., Xu, W., Yang, N., Danko, T., Chen, L., Wernig, M. & Sudhof, T. C. 2013. Rapid single-step induction of functional neurons from human pluripotent stem cells. Neuron, 78, 785–98.

